# Fine-mapping and comparative genomic analysis reveal the gene composition at the *S* and *Z* self-incompatibility loci in grasses

**DOI:** 10.1101/2022.07.18.499170

**Authors:** Marius Rohner, Chloé Manzanares, Steven Yates, Daniel Thorogood, Dario Copetti, Thomas Lübberstedt, Torben Asp, Bruno Studer

## Abstract

Self-incompatibility (SI) is a genetic mechanism of hermaphroditic plants to prevent inbreeding after self-pollination. Allogamous Poaceae species exhibit a unique gametophytic SI system controlled by two multi-allelic and independent loci, *S* and *Z.* Despite intense research efforts in the last decades, the genes that determine the initial recognition mechanism are yet to be identified. Here, we report the fine-mapping of the *Z*-locus in perennial ryegrass (*Lolium perenne* L.) and provide evidence that the pollen and stigma components are determined by two genes encoding DUF247 domain proteins (*ZDUF247-I* and *ZDUF247-II*) and the gene *sZ*, respectively. The pollen and stigma determinants are located side-by-side and were genetically linked in 10,245 individuals of two independent mapping populations segregating for *Z*. Moreover, they exhibited high allelic diversity as well as tissue-specific gene expression, matching expected characteristics of SI determinants known from other systems. Revisiting the *S*-locus using the latest high-quality whole-genome assemblies revealed a similar gene composition and structure as found for *Z*, supporting the hypothesis of a duplicated origin of the two-locus SI system of grasses. Ultimately, comparative genomic analyses across a wide range of self-compatible and self-incompatible Poaceae species revealed that the absence of a functional copy of at least one of the six putative SI determinants is accompanied by a self-compatible phenotype. Our study provides new insights into the origin and evolution of the unique gametophytic SI system in one of the largest and economically most important plant families.

## Introduction

The mating systems and mechanisms behind sexual reproduction of flowering plants are diverse: monoecious plants produce flowers with only one reproductive organ, either female or male, promoting cross-pollination (Willson 1983). Hermaphroditic plants developed various strategies promoting cross-pollination, determined, for example, by the morphology of the reproductive organs (Ganders 1979) or by differences in the maturity of these organs (Lloyd and Webb 1986).

Self-incompatibility (SI) is a mechanism preventing self-pollination upon self-pollen recognition by the female organ. Different genetic mechanisms exist in angiosperms (De Nettancourt 1977; Takayama and Isogai 2005). In most flowering plants, the recognition of self-pollen by the pistil is genetically controlled by a single multi-allelic locus, the *S*-locus. The *S*-locus encodes at least two closely linked genes, representing the male and female SI determinants (Fujii et al. 2016). The same *S*-allele specificity expressed by the pollen and the pistil will halt pollen tube development and hence, successful fertilization (Takayama and Isogai 2005).

In-depth knowledge about the underlying genetic control has been acquired for three single locus multi-allelic SI systems: the *S*-RNase type SI system (McClure et al. 1989; Kao and Tsukamoto 2004; Sijacic et al. 2004; Williams et al. 2015; Sassa 2016), the Papaveraceae type SI system (Foote et al. 1994; Wheeler et al. 2009; Poulter et al. 2010; Wilkins et al. 2014; Wang et al. 2018); and the Brassicaceae type SI system (Nasrallah et al. 1987; Schopfer et al. 1999; Takasaki et al. 2000; Sehgal and Singh 2018). The diverse identity of the genetic determinants in these well-studied SI systems strongly supports the hypothesis that different SI systems have evolved independently in different lineages (Steinbachs and Holsinger 2002; Charlesworth et al. 2005). Despite their profound differences, several evolutionary features are shared, such as the high allelic and nucleotide diversity within a species but also the low nucleotide variation between the SI determinants of the same allelic specificity (Charlesworth et al. 2005). Furthermore, the suppression of recombination between the male and female SI determinants in SI systems is considered essential, as a recombination event may produce a nonfunctional SI haplotype leading to the breakdown of SI (Fujii et al. 2016).

SI in the grass family (Poaceae) is yet to be elucidated, despite early research by Lundqvist dating back to 1954 (Lundqvist 1954). In grasses, SI is reported in many tribes such as Triticeae (*Secale cereale* L., *Hordeum bulbosum* L.), Paniceae (*Panicum virgatum* L.), Oryzeae (*Oryza longistaminata* A. Chev. & Roehr), Andropogoneae (*Miscanthus sinensis* Anderss.) and the Poeae (*Festuca pratensis* Huds., *Lolium perenne* L., *Lolium multiflorum* Lam.) (see Li et al. (1997) and Do Canto et al. (2016) for a complete list). The SI system in grasses is gametophytically controlled and genetically governed by two multi-allelic and independent loci, *S* and *Z* (Lundqvist 1954; Hayman 1956; Cornish et al. 1979). Self-recognition is based on the interaction between male and female determinants of both loci. The fertilization is halted when *S-* and *Z*-haplotypes of the pollen are matched in the stigma. The recognition of self-incompatible pollen in grasses, followed by the inhibition of the pollen tube growth, is very rapid, occurring at the stigma surface within minutes after germination (Shivanna et al. 1982). The downstream reaction upon the initial self/nonself-recognition is unknown. The involvement of calcium (Ca^2+^)-induced signaling transduction, protein phosphorylation, and the proteolysis pathway have been reported in preliminary studies (Wehling et al. 1994; Klaas et al. 2011). The current knowledge suggests that self-incompatible species of the entire Poaceae family share the same SI system, similarly as all dicotyledonous species investigated at the molecular level belonging to the same family share the same SI system (Li et al. 1997; Baumann et al. 2000). In perennial ryegrass (*L. perenne*), the *S-* and the *Z*-locus have been mapped to chromosomes 1 and 2, respectively, using genetic linkage mapping (Thorogood et al. 2002). These two loci have also been located on chromosomes 1 and 2 of rye (*S. cereale*; Wricke & Wehling, 1985; Gertz & Wricke, 1989) and sunolgrass (*Phalaris coerulescens* Desf.; Bian et al., 2004), for example. The syntenic region in self-compatible rice (*Oryza sativa* L.) can be found on chromosome 5 for *S* and chromosome 4 for *Z* (Jones et al. 2002).

More recently, the *S*-locus has been mapped to a 0.1 centimorgan (cM) region by Manzanares et al. (2016) in perennial ryegrass, containing eight genes. The gene *SDUF247* (or *LpSDUF247*, as isolated in *L. perenne*) has been suggested as the gene encoding for the pollen component, due to its high sequence diversity and the fact that the allelic sequences observed at that gene were fully predictive for the *S*-locus genotypes known to segregate in the population used for fine-mapping. Furthermore, within *SDUF247*, a frameshift mutation has been identified within self-compatible darnel (*Lolium temulentum* L.), whereas all self-incompatible species analyzed within the *Festuca-Lolium* species complex were predicted to encode functional SDUF247 proteins. However, due to the absence of a contiguous genome sequence at the *S*-locus, the identity of the female component remains elusive (Manzanares et al., 2016).

Fine-mapping of the *Z*-locus is less advanced: In rye, the genomic region containing the *Z*-locus was narrowed down to 1.5 cM on chromosome 2RL (Hackauf and Wehling 2005). Shinozuka et al. (2010) identified the orthologous region spanning 60 kb on chromosome 5 in *Brachypodium distachyon* (L.) P. Beauv.. Using a comparative genomics approach based on the synteny between chromosome 5 of *B. distachyon* and chromosome 2 of perennial ryegrass, BAC clones co-segregating with the *Z*-locus were identified and used for sequencing. From this study, a gene encoding for a protein containing a DUF247 domain has been identified in the *Z*-locus region of perennial ryegrass, as well as three other candidate genes (Shinozuka et al. 2010).

Longstamen rice (*O. longistaminata*), a self-incompatible African rice species, was recently reported to have maintained the two-locus gametophytic SI system of grasses (Lian et al. 2021). Comparative genomic analysis enabled the identification of the gene orthologous to the putative male *S*-locus determinant of perennial ryegrass (*LpSDUF247*). The gene named *OlSS1* encodes for a member of the DUF247 protein family. A second gene (*OlSS2*), also predicted to encode for a protein of the DUF247 family, was identified nearby. Sequence polymorphism analysis of the genes adjacent to *OlSS* led to the identification of a possible female determinant at *S*, *OlSP*. The *OlSP* gene contains an N-terminal YfaZ domain of unknown function, and an ortholog to this gene in *H. bulbosum (HPS10*) has been previously presented as a possible candidate for the female determinant at the *S*-locus (Kakeda et al. 2008; Kakeda 2009). The reported high density of sequence polymorphisms and expression data for the identified genes at the *S*-locus in *O. longistaminata* showed that *OlSS1* and *OlSS2* are plausible candidate genes for the male determinant, whereas *OlSP* likely encodes for the female determinant at *S* (Lian et al. 2021).

Whole-genome sequences and high-quality assemblies thereof have been established for several major self-compatible crop species within the Poaceae family, for example for rice (*O. sativa*, Stein et al. 2018), maize (*Zea mays* L., Schnable et al. 2009), barley (*Hordeum vulgare* L., Mayer et al. 2012), rye (*S.cereale* (Li et al. 2021; Rabanus-Wallace et al. 2021), wheat (*Triticum aestivum* L., Appels et al. 2018), and purple false brome (*B. distachyon*, Vogel et al. 2010). In contrast, the genomic resources available for outbreeding forage grasses like perennial ryegrass, Italian ryegrass (*L. multiflorum*), orchardgrass (*Dactylis glomerata* L.), and meadow fescue (*F. pratensis*) are limited. The primary limitations hampering the development of high-quality genome assemblies within outbreeding forage grasses are their high level of heterozygosity and the high content of repetitive sequences within the genome (Byrne et al. 2015). In recent years, more contiguous genome assemblies have become available for forage grasses and non-major crop species, including a reference-grade genome assembly of a doubled haploid genotype of perennial ryegrass (Frei et al. 2021), a high-quality draft diploid genome assembly of Italian ryegrass (Copetti et al. 2021), and a chromosome-scale diploid genome assembly of orchardgrass (Huang et al. 2020). The concurrent availability of high-quality Poaceae genome assemblies from self-incompatible and self-compatible species finally allows for an intensive comparative genomics approach to investigate the underlying genetic basis for SI.

The main objective of this study was to further characterize and advance our understanding of the two-locus gametophytic SI system in Poaceae species by identifying the male and female determinants at the *S*- and *Z*-locus. Specifically, we aimed to locate the *Z*-locus through fine-mapping in perennial ryegrass using a number of mapping individuals sufficiently high to reach gene-scale resolution. Learning from the gene composition, order, and orientation at the *Z*-locus, we further aimed to reconstruct the gene content at the *S*-locus and compare it to other species of the *Festuca-Lolium* species complex and grasses in general. Finally, through a complementary set of genetic analyses, including sequence diversity and gene expression analysis, we aimed to identify the *S*- and *Z*-locus determinants and distinguish between the male and female components of the SI system present in the family of grasses.

## Results

### Fine-mapping of the *Z*-locus in perennial ryegrass

A total of 10,245 plants from two genetically unrelated perennial ryegrass populations, hereafter referred to as VrnA-XL and DTZ, were used for fine-mapping. With a similar approach as described by Manzanares et al. (2016), the two markers CADELP and Lp02_555 flanking the *Z*-locus identified a total of 89 and 99 recombination events in VrnA-XL and DTZ, respectively (Supplementary Table 1).

To establish the physical structure at the *Z*-locus and to locate the recombination events, the perennial ryegrass BAC libraries described by Farrar et al. (2007) were screened using the marker TC116908 (Hackauf and Wehling 2005). The BAC library constructed from the genotype NV#20F1-30 (hereafter referred to as F1-30) was particularly suitable, as F1-30 is one of the two parental genotypes that was used to develop VrnA-XL. The BAC clone P205C9H17P, identified to contain the *Z*-locus of F1-30, was grown, its DNA was isolated, and sequenced. Sequence assembly reconstructed a 99,639 bp long single contig of P205C9H17P (GenBank accession number xxx), which was used to develop DNA markers for fine-mapping (Supplementary Table 1). Projection of the recombination events from the two different fine-mapping populations VrnA-XL and DTZ on P205C9H17P identified a region of 37,125 bp co-segregating with the *Z*-locus (hereafter referred to as haplotype P205), delimited by the markers BAC_BEG and 37600 (Supplementary Table 1, Figure 1).

**Figure 1:**
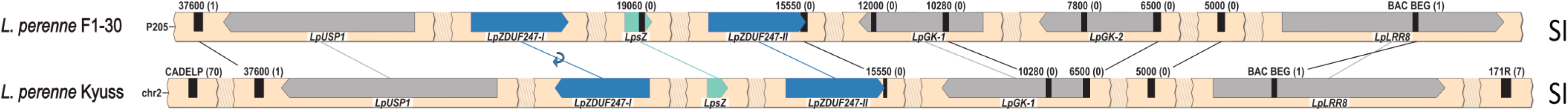
The gene composition of the genome region co-segregating with the *Z*-locus in perennial ryegrass (*Lolium perenne* L.). Given are the genotypes F1-30 (haplotype P205, above) and in the doubled haploid genotype Kyuss (below). The sequence of the *Z*-locus is continuous, but for clarity, the gene- and marker-less regions are represented as shaded breaks. The genes are represented with bars, and their orientation is shown with the pointy side representing the 3’ end. The self-incompatibility candidate genes are colored in teal and blue. The markers used for the fine-mapping are represented by black bars, and the number of recombinants between each marker is indicated between brackets. The synteny between homolog genes is illustrated with lines connecting the two haplotypes, and in case of orientation change, a small circular arrow is used. On the right, the compatibility phenotype is indicated (SI = self-incoffmpatible).

The annotation of the genome region co-segregating with the *Z*-locus was done using available genomic resources (Byrne et al. 2015; Begheyn et al. 2018; Copetti et al. 2021; Frei et al. 2021), gene prediction software (Stanke and Morgenstern 2005), and a manual BLAST-based approach. Six genes have been identified on P205: *LpUSP1, LpZDUF247-I, Lolium perenne* stigma Z (*LpsZ), LpZDUF247-II, LpGK*, and *LpLRR8* (Figure 1 and Table 1). The *Z*-locus as revealed for P205 was compared to the reference-grade perennial ryegrass genome of the doubled haploid genotype Kyuss (Frei et al. 2021): While the overall gene order was conserved between the two perennial ryegrass haplotypes, the partial duplication of the *LpGK* was missing in Kyuss. Furthermore, the orientation of *LpZDUF247-I* was not conserved between the two perennial ryegrass genotypes (Figure 1). Two *Z*-locus genes containing a DUF247 domain are present in both genotypes and have been annotated as two different genes (*LpZDUF247-I* and *LpZDUF247-II*), their nucleotide sequence being too different to be considered as a recent gene duplication.

**Table 1:**
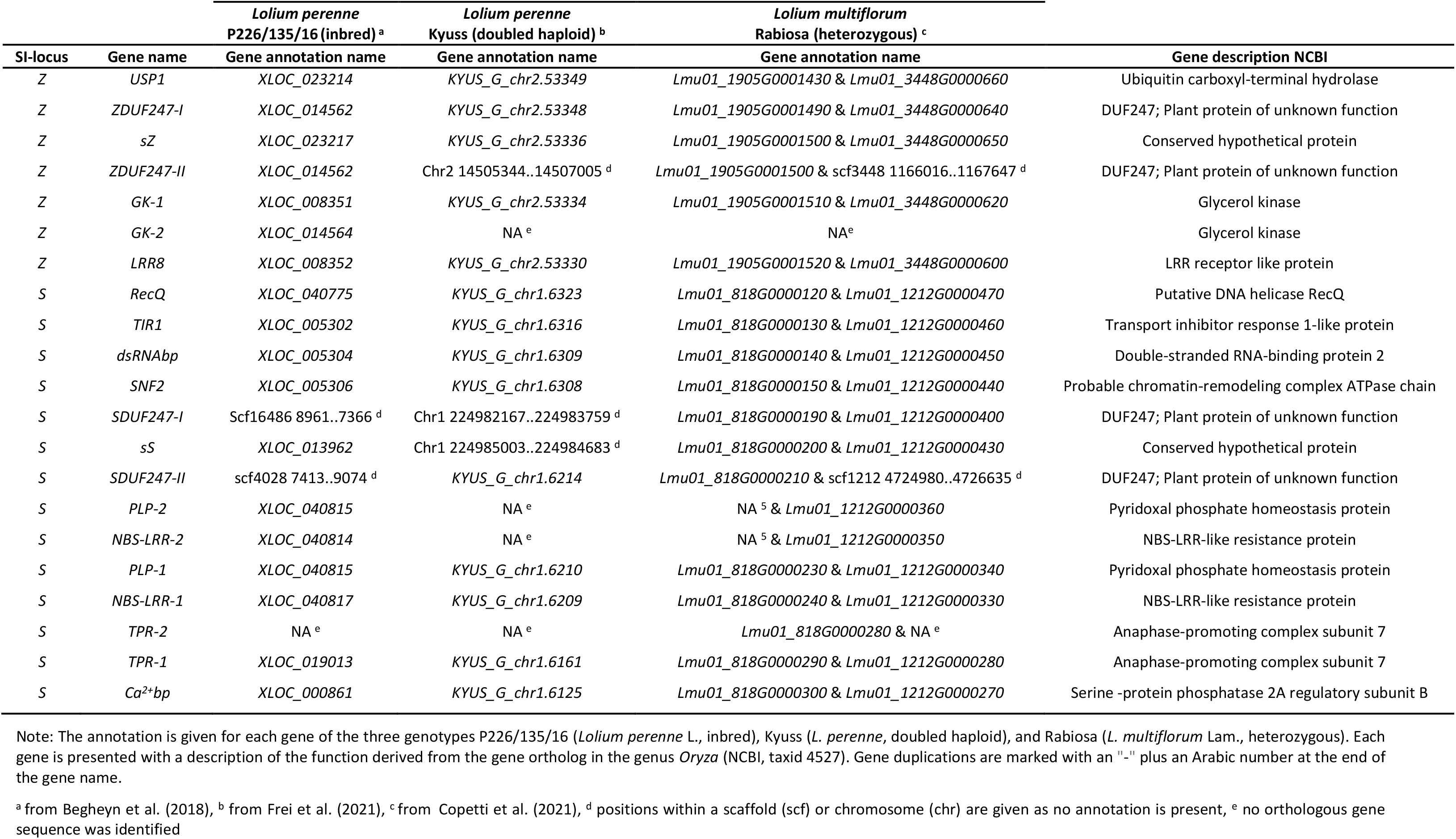
Gene composition at the *Z*- and the *S*-locus of the gametophytic self-incompatibility system of grasses.

### Comparative genomics – Synteny of the *S*- and the *Z*-locus in the Poeae tribe and the Poaceae family

The *S*-locus in perennial ryegrass, as described by Manzanares et al. (2016), contained nine unique genes, and the putative male determinant was identified as a gene harboring a DUF247 domain (*LpSDUF247*, hereafter referred to as *LpSDUF247-I*). In order to establish a contiguous genome sequence covering the *S*-locus, thereby identifying genes potentially missed in the fragmented assembly used by Manzanares et al. (2016), a comparative genomics analysis with the available genome sequence resources of *Lolium* species (Table 2) was applied. By such analysis, three additional genes were found: *LpTPR*, another gene encoding for a DUF247 domain-containing protein (*LpSDUF247-II*), and *Lolium perenne* stigma *S (LpsS*) (Table 1).

**Table 2:**
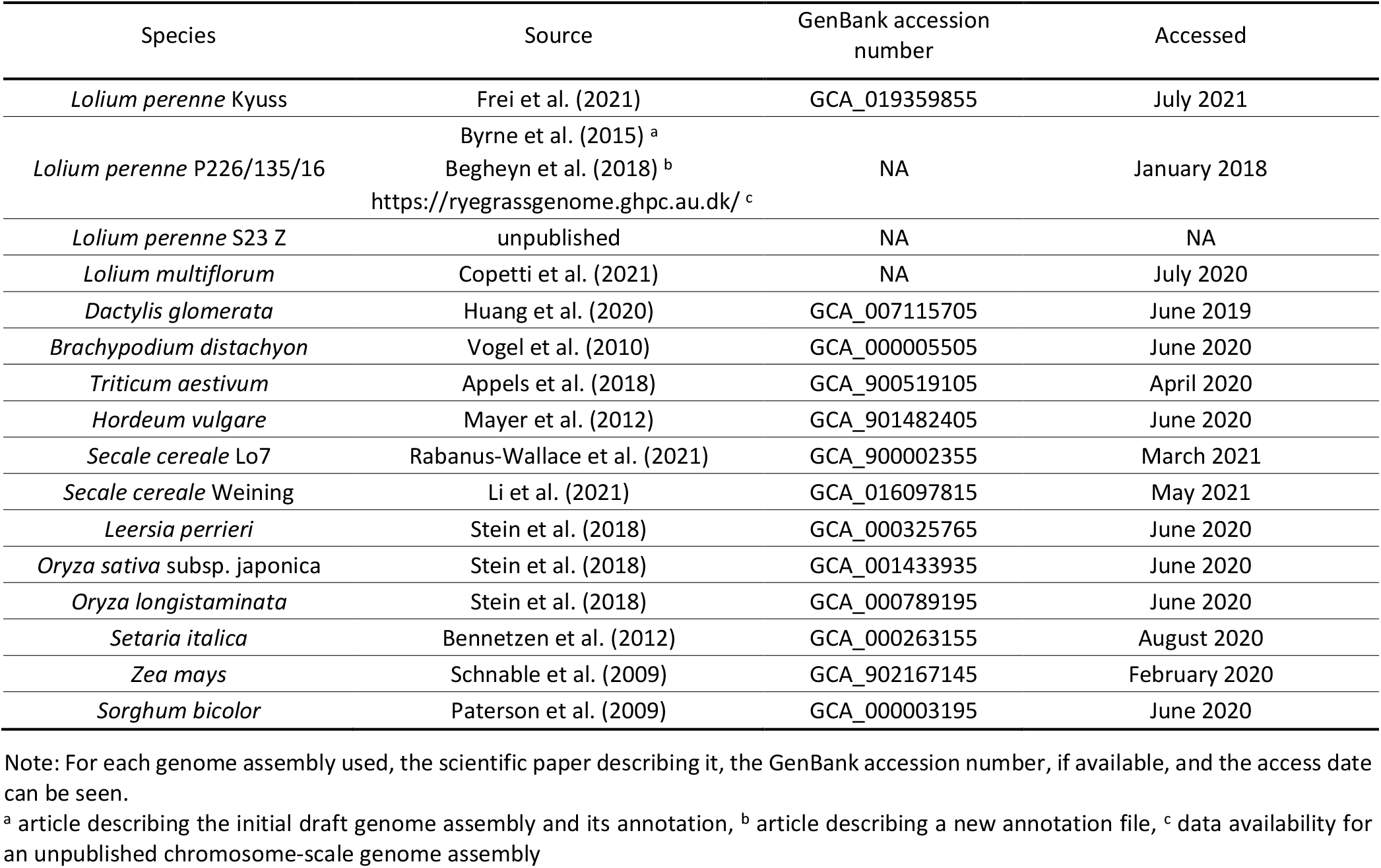
Genome data used for the comparative genome analysis.

Comparison of the structure and predicted function of the genes found at the *S*-locus with the six newly identified genes at the *Z*-locus revealed similarities between the two SI loci in *Lolium* species (Table 1). The *SI-DUF247* genes (*SDUF247-I, SDUF247-II, ZDUF247-I*, and *ZDUF247-II*) as well as *sS*, and *sZ* are here referred to as SI candidates, based on the already identified male determinant (*SDUF247-I*) by Manzanares et al. (2016) and the potential duplicative origin of the two-locus SI system in grasses (Lundqvist 1962).

The *SI-DUF247* genes found within the *L. perenne* genotype P226/135/16, the *L. perenne* genotype Kyuss, and the *L. multiflorum* genotype Rabiosa all belong to the same protein family DUF247 (pfam03140), have a similar protein size of 507-559 amino acids, and display the same intron-less gene structure. The domain hits using InterProScan (Blum et al. 2021) predicted a non-cytoplasmic domain at the C-terminus, followed by a transmembrane domain and a small cytoplasmic domain at the N-terminus for all SI-DUF247s analyzed.

The genes *sS* and *sZ* extracted from the same three genotypes displayed a conserved structure containing two exons. The translated protein sequences were 82 to 99 amino acids long and harbored a signal peptide at the N-terminus followed by a non-cytoplasmic domain.

To further study the gene content, order, and orientation at the *S*- and the *Z*-locus, the comparative genomic analysis was extended to include a wide range of self-compatible and self-incompatible species belonging to the tribe Poeae (Figure 2 and Figure 3) and to the family Poaceae (Figure 4 and Figure 5), as summarized in Table 2.

**Figure 2:**
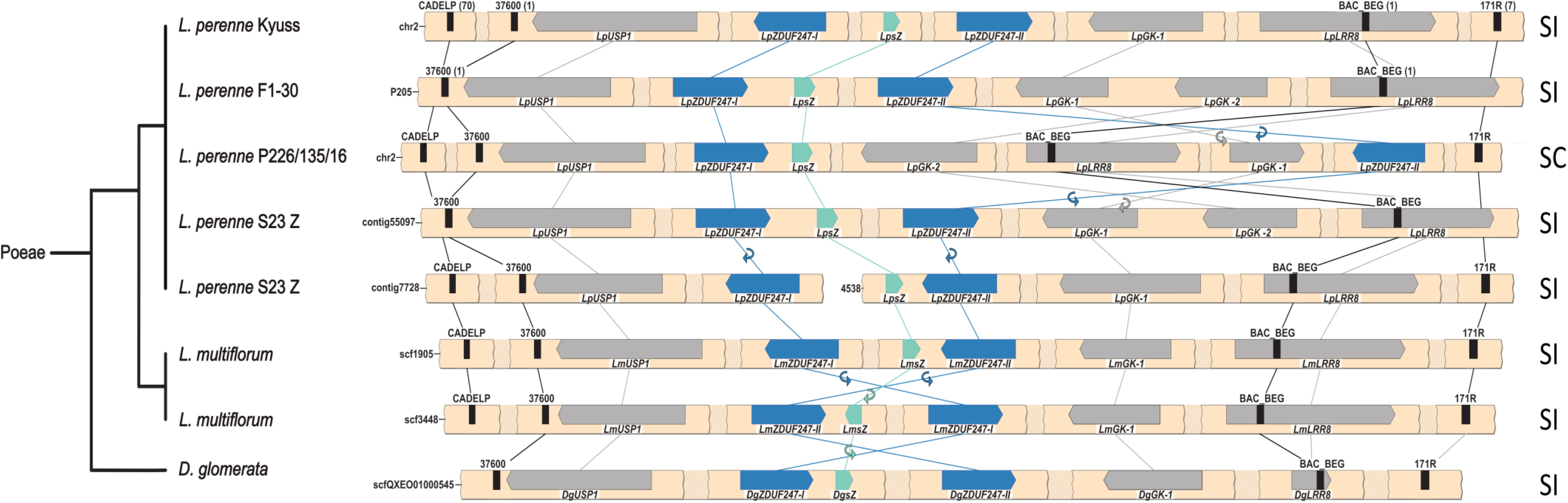
Synteny maps of the *Z*-locus of multiple genotypes from the Poeae tribe. *Lolium perenne* L. genotype S23 Z and *Lolium multiflorum* Lam. represent diploid assemblies; therefore, both haplotypes are represented. The phylogenetic tree (left) representing the different species was drawn according to the NCBI taxonomy database. The gene- and marker-less regions are represented as shaded breaks, and a white space indicates an assembly gap. The genes present at the *Z*-locus are represented with directed arrows, and self-incompatibility candidate genes are colored in teal and blue. The gene orientation is shown with the pointy side representing the 3’ end. The markers used for the fine-mapping are represented by black bars, and the number of recombinants for each marker is indicated between brackets for *L. perenne* Kyuss and F1-30. The synteny between genes is illustrated by lines, and in case of orientation change, a small circular arrow is used. In addition, the compatibility phenotype of the genotype is indicated on the right: self-incompatible (SI) or self-compatible (SC).

**Figure 3:**
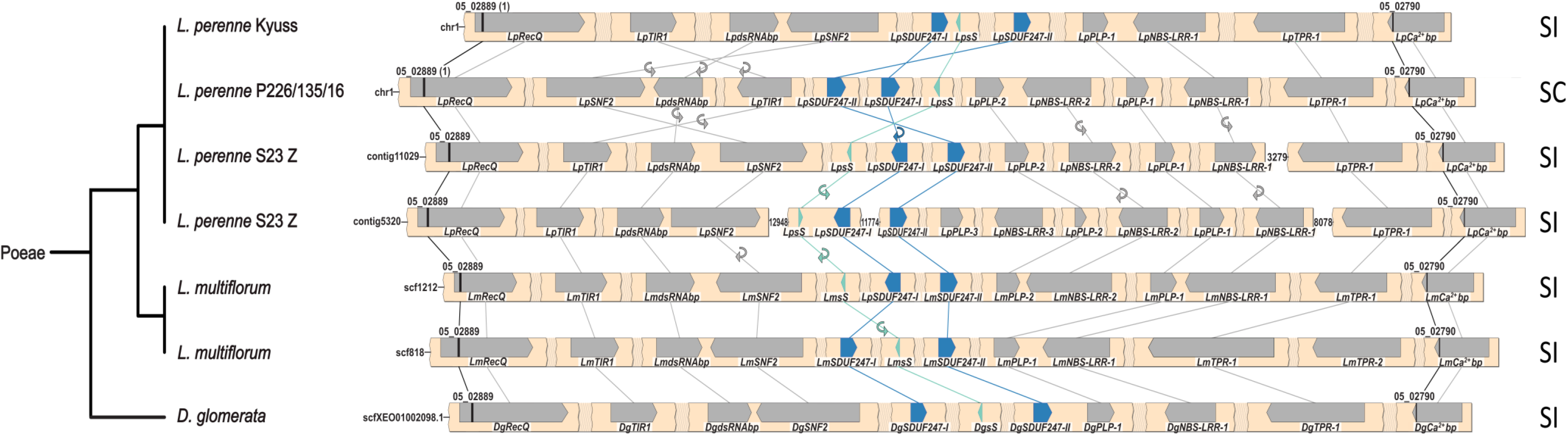
Synteny maps of the *S*-locus of multiple genotypes from the Poeae tribe. *Lolium perenne* L. genotype S23 Z and *Lolium multiflorum* Lam. represent diploid assemblies; therefore, both haplotypes are represented. The phylogenetic tree (left) representing the different species was drawn according to the NCBI taxonomy database. The gene- and marker-less regions are represented as shaded breaks, and a white space indicates an assembly gap. The genes present at the *S*-locus are represented with directed arrows and the self-incompatibility candidate genes are colored in teal and blue. The gene orientation is shown with the pointy side representing the 3’ end. The markers used for the fine-mapping are represented by black bars, and the number of recombinants for each marker is indicated between brackets for *L. perenne* Kyuss and P226/135/16. The synteny between genes is illustrated by lines, and in case of orientation change, a small circular arrow is used. In addition, the compatibility phenotype of the genotype is indicated on the right: self-incompatible (SI) or self-compatible (SC).

**Figure 4:**
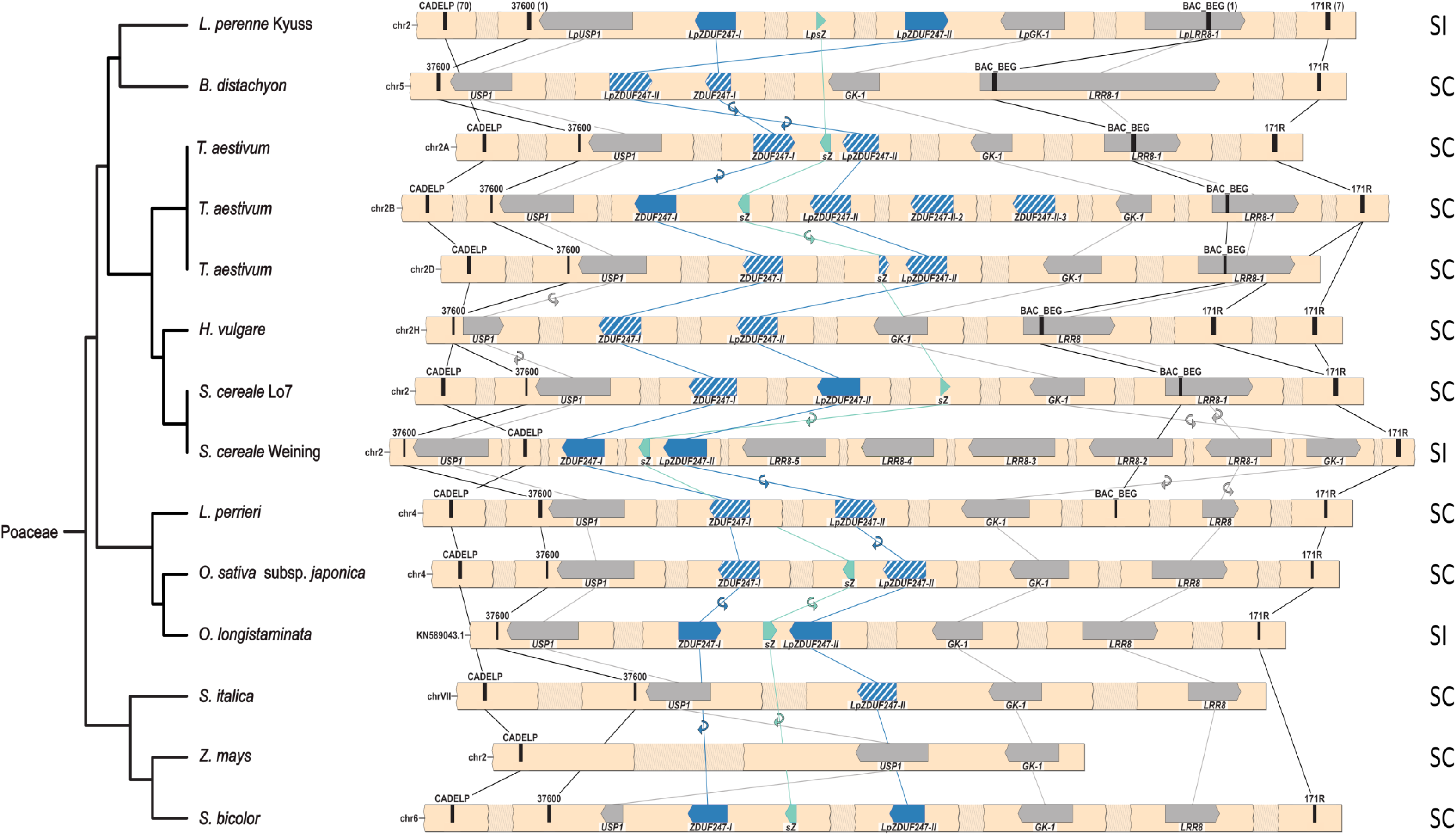
Synteny maps of the *Z*-locus of eleven Poaceae species. The phylogenetic tree (left) representing the different species was drawn according to the NCBI taxonomy database. *Triticum aestivum* L. represents an allohexaploid composed of homologous genomes (A, B, and D). The gene- and marker-less regions are represented as shaded breaks, and a white space indicates an assembly gap. The genes present at the *Z*-locus are represented with directed arrows and the self-incompatibility candidate genes are colored in teal and blue. A nonfunctional gene copy of the self-incompatibility candidates is indicated with a white striped pattern. The gene orientation is shown with the pointy side representing the 3’ end. The markers used for the fine-mapping are represented by black bars, and the number of recombinants for each marker is indicated between brackets for *L. perenne* Kyuss. The synteny between genes is illustrated by lines, and in case of orientation change, a small circular arrow is used. In addition, the compatibility phenotype of the genotype is indicated on the right: self-incompatible (SI) or self-compatible (SC).

**Figure 5:**
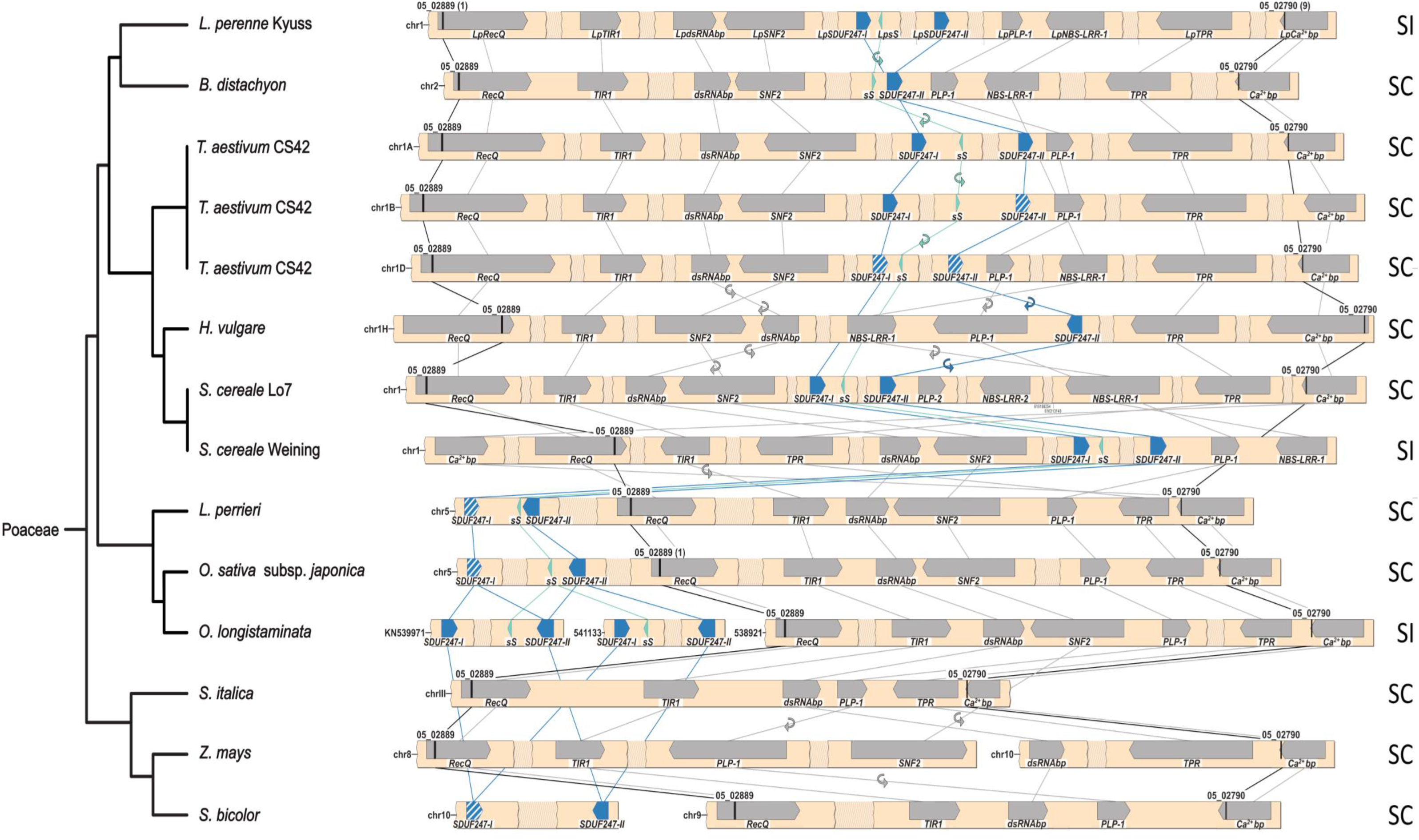
Synteny maps of the *S*-locus of eleven Poaceae species. The phylogenetic tree (left) representing the different species was drawn according to the NCBI taxonomy database. *Triticum aestivum* L. represents an allohexaploid composed of three homologous genomes (A, B, and D). The gene- and marker-less regions are represented as shaded breaks, and a white space indicates an assembly gap. The genes present at the *S*-locus are represented with directed arrows, and the self-incompatibility candidate genes are colored in teal and blue. A nonfunctional gene copy of the self-incompatibility candidates is indicated with a white striped pattern. The gene orientation is shown with the pointy side representing the 3’ end. The markers used for the fine-mapping are represented by black bars, and the number of recombinants for each marker is indicated between brackets for *L. perenne* Kyuss. Lines illustrate the synteny between genes, and in case of orientation change, a small circular arrow is used. In addition, the compatibility phenotype of the genotype is indicated on the right: self-incompatible (SI) or self-compatible (SC).

Generally, a high level of genome synteny was observed at the *S*- and the *Z*-locus. The highest degree of synteny was found within species and genotypes from the Poeae tribe. Minor deviations included changes in the gene orientation, the duplication level of certain genes, and the gene order of the SI candidate genes (Figure 2 and Figure 3). In contradiction to the high degree of synteny within the Poeae tribe stands the draft genome assembly of the self-compatible *L. perenne* genotype P226/135/16, which displayed a unique gene order at both the *S*- and the *Z*-locus (Figure 2 and Figure 3). At the *Z*-locus, the region harboring *LpGK-1* and *LpZDUF247-II* was inverted and reintegrated. At the *S*-locus, the region harboring *LpSNF2, LpdsRNAbp*, and *LpTIR* was also inverted.

The *S*- and the *Z*-locus identified in genotypes outside the Poeae tribe showed mainly a high synteny with the *S*- and the *Z*-locus of *Lolium* spp. (Figure 4 and Figure 5), especially closely related species of the Triticaceae tribe (e.g., *T. aestivum, H. vulgare*, and *S. cereale*). Notable gene order alterations within the Triticaceae tribe were found in the *S. cereale* genotype Weining at the *S*-locus (Figure 5). A lower but comparable degree of synteny was observed within the Oryzeae tribe (e.g., *L. perrieri*, and *O. sativa* subsp. Japonica) and *Sorghum bicolor* L., except that the SI candidate genes are located outside of the perennial ryegrass *S*-locus. The gene cluster consisting of *SDUF247-I, SDUF247-II*, and *sS* was not flanked by the perennial ryegrass flanking markers or the flanking genes. For *O. sativa* subsp. *Japonica*, the SI candidate genes were 3.14 Mbp upstream of the flanking marker 05_02889 (Manzanares et al. 2016). In *L. perrieri*, the distance was 2.1 Mbp between the flanking marker 05_02889 and the SI candidate genes. For *O. longistaminata*, the SI candidate genes at S were present as duplication on individual scaffolds. However, whether the two copies result from a duplication or if both *S*-haplotypes were included in the haploid assembly remains elusive. In *S. bicolor*, the SI candidate genes were located on chromosome 10, whereas the *S*-locus flanking markers and flanking genes were localized on chromosome 8. In *S. italica* and *Z. mays*, almost no synteny could be observed at both loci, mainly through the absence of the SI candidate genes (Figure 4 and Figure 5).

The functionality of the orthologous SI candidate genes was evaluated in addition to the synteny within the Poeae tribe and Poaceae family. A gene was considered to be functional if an open reading frame was present, leading to a protein of similar size containing the same protein motifs prediction as the SI candidate genes identified within the *Lolium* species. In the Poeae tribe, all SI candidate genes were present and assessed to be functional (Figure 6). Within the Poaceae family, orthologous genes of *sS* and *sZ* were mainly predicted to be functional, unlike most of the *SI-DUF247s* (Figure 6). Furthermore, all Poaceae species and genotypes investigated displaying a self-incompatible phenotype always harbored six functional SI candidate genes (Figure 6).

**Figure 6:**
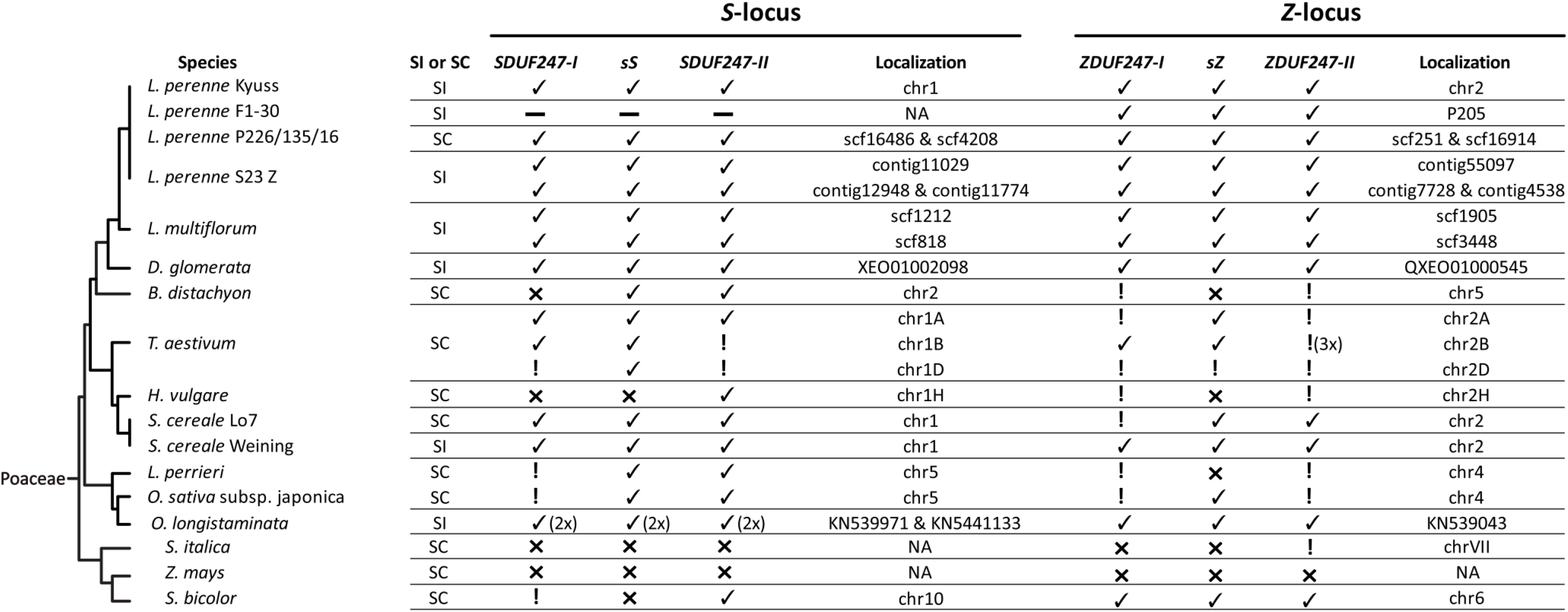
Composition of the self-incompatibility candidate genes in 17 genotypes representing 13 different Poaceae species. A phylogenetic tree was drawn according to the NCBI taxonomy database on the figure’s left. The compatibility phenotypes are indicated for each genotype: self-incompatible (SI) or self-compatible (SC). A checkmark (✓) represents the presence of a functional gene, an exclamation mark (!) indicates that the sequence is present but was evaluated to be nonfunctional. A cross (x) means an orthologous sequence was not found. The minus sign (—) indicates that the sequence data was not available, and therefore the presence/absence of the gene cannot be determined. In addition, the position on chromosome or scaffold level of the self-incompatibility candidate genes in the genome is given. *Lolium perenne* L. genotype S23 Z and *L multiflorum* Lam. represent diploid assemblies; therefore, both haplotypes are displayed. *Triticum aestivum* L. represents an allohexaploid species leading to a triplication of the *S*- and *Z*-locus. Besides on chromosome 2B (chr2B), a nonfunctional copy of the *ZDUF247-II* was present three times. In *Oryza longistaminata* A. Chev. & Roehr, the gene copies of functional *S* self-incompatibility candidate genes are present on two different scaffolds.

### Phylogenetic analysis of genes located within the *S*- and the *Z*-locus

The allelic richness of the genes within the *S*- and the *Z*-locus in *Lolium* spp. was evaluated using the coding sequences from the *L. perenne* genotypes S23 Z, Kyuss, P226/135/16, F1-30, and the *L. multiflorum* genotype Rabiosa. A phylogenetic tree was constructed for each gene using the alleles present (Figure 7).

**Figure 7:**
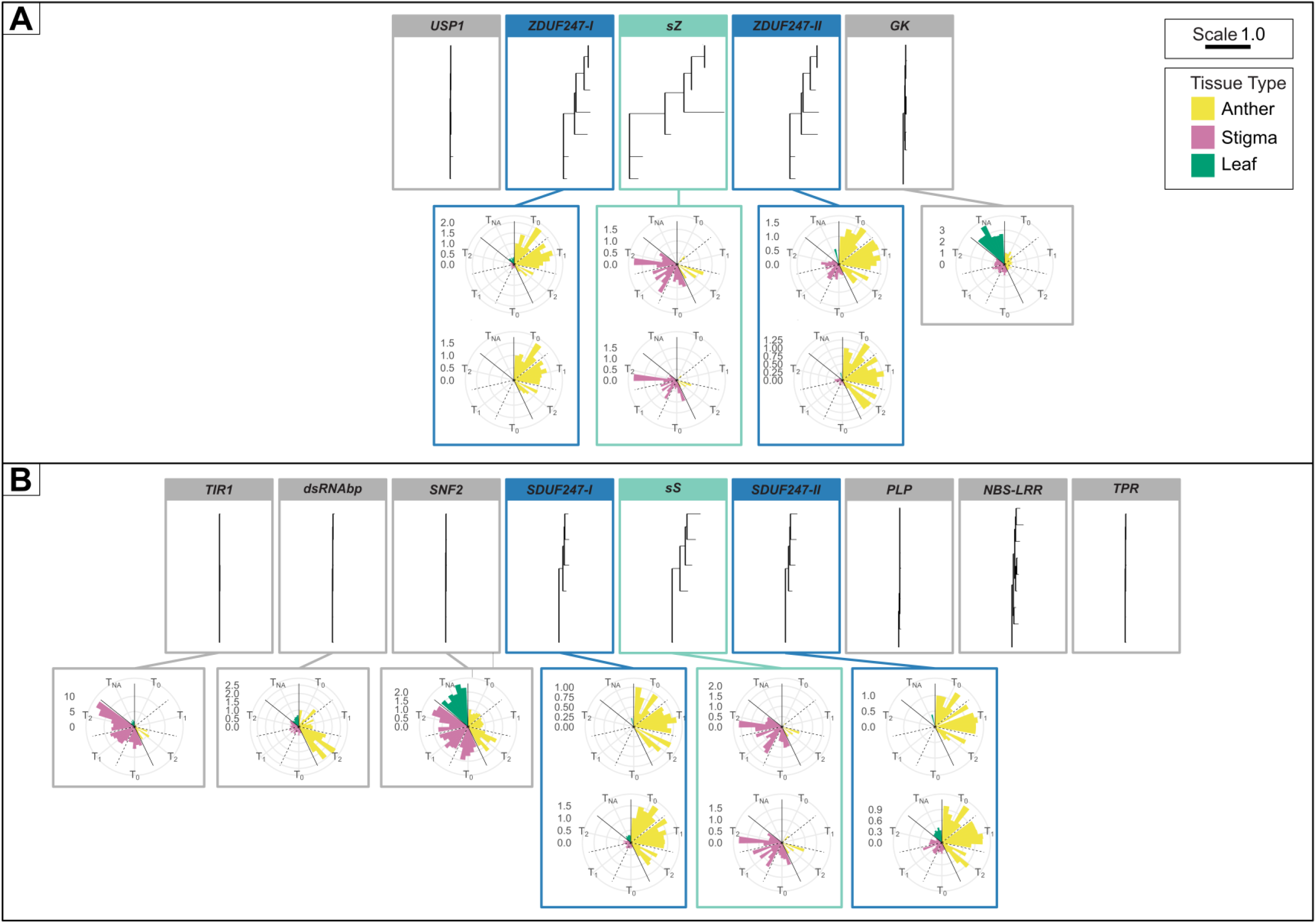
Phylogenetic trees and relative expression ratio of the genes at the *Z*-locus (A) and the *S*-locus (B). The phylogenetic trees and relative expression ratios are ordered according to the physical gene order as seen in *Lolium perenne* L. genotype Kyuss, and gene expression data is from the genotype S23 Z. The flanking genes are boxed in grey, whereas the self-incompatibility candidate genes are boxed in teal and blue. The scale bar in the top right corner represents one amino acid change per site, and the legend below shows the color code for the three tissue types used in the expression pattern analysis. For the *LpsS, LpsZ*, and the *SI-DUF247s*, the relative expression ratio measurement was explicitly performed for each allele present in the genotype. The two different alleles are displayed stacked on top of each other. For the remaining *S*- and *Z*-locus genes, the expression pattern was investigated with primers amplifying both alleles, and therefore only one polar plot is presented per gene. Only data points were included, where the C_t_ difference was below 0.5 and the percent deviation was below 3% between the two technical replicates. Therefore, the standard error is not displayed in the figure.

The SI candidate genes, as well as *NBS-LRR and LRR8*, exhibited a high allelic richness. The remaining genes at the *S*- and the *Z*-locus were highly conserved within the genus *Lolium*. To further investigate the sequence diversity of the SI candidate genes, a pairwise comparison based on a T-Coffee alignment of the amino acid sequence was performed and displayed in a heat map (Supplementary Figure 1). The sS alleles showed a mean protein sequence identity of 52.9% with a standard deviation (σ) of 6.8. The mean protein sequence identity of the sZ alleles was significantly lower, being 32.5% (σ = 9.5). The mean protein sequence identity between the sS and sZ alleles is 28.1% (σ = 4.3). The SI-DUF247s show a higher level of protein sequence conservation within the different alleles with the mean protein sequence identity, being 79.5% (σ = 1.9) for SDUF247-I, 75.2 % (σ = 3.2) for SDUF247-II, 60% (σ = 6.7) for ZDUF247-I, and 53% (σ = 6.9) for ZDUF247-II. A comparison between all paralogues and genotypes of the SI-DUF247s revealed mean protein sequence identities ranging from 40.9% to 45.2%.

### Expression analysis of genes located within the *S*- and the *Z*-locus

To identify the female and the male components involved in the SI reaction, the expression pattern of the genes within the *S*- and the *Z*-locus in *L. perenne* were analyzed using RT-qPCR. The expression pattern was investigated for the six SI candidate genes, one flanking gene at the *Z*-locus (*LpGK*) and three at *S*-locus (*LpTIR1*, *LpdsRNAbp* and *LpSNF2*). Samples from *L. perenne* genotype S23 Z were taken from anther and stigma tissue at three development stages: one week before flowering (time point 0), two to three days before flowering (time point 1), and the day of flowering (time point 2). In addition, leaf tissue of S23 Z was sampled with no specific time point (time point NA).

The high allelic diversity of *LpsS, LpsZ* and the *LpSI-DUF247s* made it necessary to analyze both alleles for these genes individually. The expression data were visualized in a polar chart (Figure 7), and a scatter plot (Supplementary Figure 2) as the relative gene expression ratio calculated according to Pfaffl (2001). Moreover, the ΔC_t_ values (C_t_ value of the gene of interest minus the geometrical mean C_t_ value of the reference genes) were calculated to allow the comparison of the expression levels of different genes within the same sample and are displayed in a heat map (Supplementary Figure 3).

Polar plots of the relative gene expression show that the *LpSI-DUF247s* genes display a tendency of anther-specific expression with decreasing expression towards the day of flowering. In leaf and stigma tissue, little or no *LpSDUF247* expression was measured (Figure 7, Supplementary Figure 2, and Supplementary Figure 3). In contrast, the *LpsS* and *LpsZ* display a stigma-specific expression pattern. However, high expression was also measured in anther tissue for three biological replicates (A, C, and F) at time point 2 and the biological replicate E at time point 1 (Supplementary Figure 2 and Supplementary Figure 3). Furthermore, *LpsZ* and *LpsS* in stigma tissue displayed the highest expression levels relative to the reference genes (Supplementary Figure 3)*. LpGK* and *LpSNF2* did not display a tissue-specific pattern and were expressed in all tissue types and development stages, with *LpGK* being overexpressed in leaves. The *LpdsRNAbp* showed an anther-specific expression with an apparent upregulation on the day of flowering. *LpTIR1* displayed stigma-specific expression according to the RT-qPCR experiment. *LpTIR1* showed up to a 14 times higher expression in the stigma on the day of flowering compared to the control sample (Anther T_1_ biological replicate A).

## Discussion

After almost 70 years of research on the two-locus gametophytic SI system of grasses, we established the gene content, order, and composition at the *S*- and the *Z*-locus. For each locus, two male and one female determinant are suggested to govern the SI system in grasses. All four putative male determinants have a similar gene structure and encode for proteins belonging to the same family (DUF247). The putative female determinants at *S* (*sS*) and *Z (sZ*) are also of similar gene structures and are predicted to code for secreted proteins with no known family membership. Typical characteristics of SI determinants could be observed for the putative SI genes, including genetic and physical linkage, high allelic richness, high sequence diversity, and an anther- or stigma-specific expression pattern. Furthermore, the absence of a functional copy of at least one of the six putative SI determinants is accompanied by a self-compatible phenotype within the Poaceae species.

According to the hypothesis of Lundqvist (1962), the two-locus SI system in grasses originated from a duplication of a one-locus SI system. Following this hypothesis, the male and female determinants at *S* and *Z* represent gene duplicates with a similar gene sequence and structure (Yang et al. 2008). The *sZ*, *ZDUF247-I*, and *ZDUF247-II* at the *Z*-locus are the only three genes for which genes of similar structure and sequence could be found at the *S*-locus (*sS*, *SDUF247-I*, and *SDUF247-II*) within self-incompatible grass species. The same protein family membership (DUF247), the same *in silico* motif predictions, similar protein size, and their conserved intron-less gene structures are clear indicators of a shared origin for the *SI-DUF247s* genes within grasses. The presence of two *SI-DUF247s* genes at each locus also suggests that a duplication event within a SI locus occurred prior to the duplication of the whole locus. For the *sS* and *sZ*, the data also indicates a duplicative origin as both are of similar size, have the same gene structure (one intron), and have the same *in silico* protein motif prediction within self-incompatible grass species. Furthermore, the presence of one additional coding sequence similar to *SDUF247-II* outside of the *S*- and *Z*-locus indicates even a further duplication event. The additional *SDUF247-II* sequence is located on chromosome 6 on the reference-grade *L. perenne* genome assembly without an annotation (position: 11313479-11315143).

The putative SI determinants in grasses were genetically and physically linked, indicating that they are inherited as a unit. The inheritance as a unit of the SI determinant is necessary as recombination events between SI determinants may lead to a breakdown of the SI system as a new SI-haplotype is generated, consisting of SI determinants expressing different SI specificities (Takayama and Isogai 2005). However, at both loci in self-incompatible grass species, the gene order and orientation vary, indicating preceding recombination events within the *S*- and *Z*-locus that, interestingly, did not lead to a breakdown of the SI system. Whereas unlikely due to the high quality of the genome assemblies used in this study, assembly errors at *S* and *Z* would represent an alternative explanation for the observed gene order and orientation changes.

Besides the unifying scheme that the SI determinants must be inherited as one segregating unit, the physiological reaction and SI genetics dictate that male determinants must be expressed in the pollen. In contrast, the female determinants must be expressed in the stigma (Takayama and Isogai 2005). The putative SI determinants, already being in line with the duplicative origin of the SI system in grasses and showing a close linkage, all displayed either an anther- or stigma-specific expression pattern. The *LpSI-DUF247s* having an anther-specific expression, and the *LpsS* and *LpsZ* a stigma-specific expression, allowing the conclusion that the *SI-DUF247s* represent the putative male SI determinants, whereas the *sS* and *sZ* represent the putative female determinant of the grasses SI system.

The *LpdsRNAbp* also showed anther-specific expression, whereas the *LpTIR1* showed a stigma-specific expression. Nonetheless, their direct involvement in the self-recognition process of the SI system in grasses can be excluded (*LpTIR1*) or is highly unlikely (*LpdsRNAbp*). The design of the mapping population used by Manzanares et al. (2016) for the fine-mapping of the *S*-locus dictated the presence of five alleles, and only three alleles were found for *LpTIR1*. Our analysis aligns with these findings as *TIR1* in the *Lolium* spp. analyzed showed high sequence conservation, uncharacteristic for an SI determinant. The same holds for the *dsRNAbp*, which, like *LpTIR1*, did not show a high allelic richness and sequence diversity within *Lolium* spp. In addition, the argument can be brought forward that, for *dsRNAbp* and *TIR1*, no gene with a similar gene structure or sequence is present at both loci, contradicting the duplicative origin hypothesis of the two-locus SI system in grasses.

Besides the putative SI determinants being the only genes at the *S*- and the *Z*-locus suggesting a duplicative origin and at the same time showing an anther- or stigma-specific expression, they also showed the typical evolutionary characteristics of an SI determinant, i.e., a high protein sequence diversity and a high allelic richness (Charlesworth et al. 2005). The observed protein sequence diversity of the putative SI determinants in *Lolium* spp. was comparable to the ones observed within the *S*-RNase type SI system (Ioerger et al. 1990; Ushijima et al. 1998; Williams et al. 2014; Dzidzienyo et al. 2016), the Papaveraceae type SI system (Paape et al. 2011), and the Brassicaceae type SI system (Jany et al. 2019). The sequence identities were matched best with the *S*-RNase type SI system with a highly diverse female determinant (*S*-RNase) (Ioerger et al. 1990; Ushijima et al. 1998; Dzidzienyo et al. 2016) and more conserved male determinants (*SLF*) (Williams et al. 2014). This was similar for the putative SI determinants in grasses where the putative male determinants (*SI-DUF247s*) were more conserved than the female determinants (*sS* and *sZ*).

In addition to the high sequence diversity, the expected high allelic richness was also matched. Each allelic sequence of the putative SI determinants represents a unique allele with one exception at the *S*-locus and one exception at the *Z*-locus. The sequence diversity and allelic richness analysis were limited by the sequence data being restricted to seven *Z*-loci and six *S*-loci from *Lolium* spp.. A more representative picture can be seen when the data presented here is combined with additional sequence data available. Especially for the *LpSDUF247-I*, a total of 24 allele sequences could be identified, which showed a mean protein sequence identity of 78.5% (σ = 3.7) when our data was combined with the allelic sequences identified by Manzanares et al. (2016) and Veeckman et al. (2019).

Two more genes co-segregating with the *S*- or *Z*-locus would fulfill the requirement of high allelic richness and high sequence diversity: the *NBS-LRR (NBS-LRR-1, NBS-LRR-2* and *NBS-LRR-3*) and the *LRR8*. Nonetheless, a possible SI determinant role is excluded for both. The involvement of the *NBS-LRR* in SI as an SI determinant was excluded in Manzanares et al. (2016) because the gene expression profile did not show a tissue-specific expression. Further, the *NBS-LRR* was present as gene duplication or triplication within multiple *S*-loci, and the sequences were pooled for the sequence diversity and the allelic richness analysis, biasing the results. The *LRR8* is excluded as a possible SI determinant at the *Z*-locus because a marker with one recombination (BAC_BEG) lies within the gene’s coding sequence. Further, for *LRR8* or *NBS-LRR*, a gene with a similar gene structure and sequence is not present in both loci, contradicting the hypothesis of a duplicative origin of the two-locus SI system for grasses. Possible involvement in disease resistance was predicted for both genes, representing an alternative reason for the high allelic richness and the high sequence diversity observed (Mondragón-Palomino et al. 2002; McHale et al. 2006; Ng and Xavier 2011).

As additional evidence that the here reported putative SI determinants indeed govern the SI system in grasses, a distinctive genotypic pattern within the *S*- and *Z*-locus of self-incompatible and self-compatible genotypes can be seen. All self-incompatible grass genotypes have a functional copy of the *sS*, *SDUF247-I, SDUF247-II* at *S* and *sZ*, *ZDUF247-I*, and *ZDUF247-II* at *Z*. A genotype missing a functional copy of any of the putative SI determinants shows a self-compatible phenotype. The two *S. cereale* genotypes are worth mentioning as the genotype Weining displayed a self-incompatible phenotype, and the inbred line Lo7 displayed a self-compatible phenotype. They differ genotypically as the inbred line Lo7 did not harbor a functional copy of the *ZDUF247-I*, representing an explanation for the breakdown of the SI system. In contrast, it cannot be concluded from a functional set of all six putative SI determinants that the plants show a self-incompatible phenotype. The absence of a functional gene copy of an SI determinant, disrupting the initial self-recognition process, is not the only source of self-compatibility (Do Canto et al. 2016). Similarly, a recombination event between the SI determinants, silencing of the SI determinants, or a mutation interfering with the downstream cascade of SI unlinked to the *S*- and the *Z*-locus represent other sources of self-compatibility (Do Canto et al. 2016; Cropano et al. 2021).

In our analysis, the *L. perenne* genotype P226/135/16 represented the only case where six functional SI determinants were present but no self-incompatible phenotype was reported. The source of self-compatibility remains unknown. But the loss of close linkage of the putative SI determinants at the *Z*-locus indicates a recombination event and a possible self-compatibility source (Takayama and Isogai 2005).

Based on the physiological observations of the pollen tube growth and its halt in self-incompatible grass species, Heslop-Harrison and Heslop-Harrison (1982) suggested that the male determinants must be anchored in the membrane of the pollen and that the female determinant is a secreted and diffusible protein. A similar mechanism of the SI system was also suggested by Wehling et al. (1994). The putative male determinants (SI-DUF247s) being predicted to be membrane-bound proteins and the putative female determinants (sS and sZ) predicted to be secreted into the extracellular space would agree with the suggested physiological mechanisms.

Whereas our analysis was mainly focused on *Lolium* spp., it is commonly believed that the outbreeding nature of grass species can be attributed to the same SI system (Li et al. 1997; Baumann et al. 2000). This belief is further supported by the high synteny observed of the *S*- and *Z*-locus and especially the presence of functional copies of the putative SI determinants in self-incompatible species belonging to the Poeae tribe (*L. perenne, L. multiflorum*, and *D. glomerata*), Triticeae tribe (*S. cereale*), and Oryzeae tribe (*O. longistaminata*). Furthermore, for another member of the Triticaceae tribe (*H. bulbosum*), the gene *HPS10* was presented as a possible candidate for the female determinant at the *S*-locus (Kakeda et al. 2008; Kakeda 2009). The *HPS10* is orthologous to the presented putative female determinant at the *S*-locus (*sS*). Our findings further support those of Lian et al. (2021), who reported two male candidates at the *S*-locus, *OlSS1* and *OlSS2*, orthologous towards the *SDUF247-I* and *SDUF247-II*, and one female candidate, the *OlSP*, orthologous to the *sS*.

In conclusion, our study provides multiple lines of evidence that the *SI-DUF247* are the male SI determinants in grasses at both loci (*S* and *Z*), whereas *sS* is the female determinant at *S*, and *sZ is* the female determinant at *Z*. The identification of the SI determinants enables the prediction of pollen compatibility and pollination efficiency as well as the targeted induction and exploitation of loss-of-function mutations at *S* or *Z*, leading to self-compatibility, both a quantum leap in the breeding of allogamous grass species. More broadly, our study offers new insights into the origin and evolution of the unique gametophytic SI system in one of the largest and economically most important plant family.

## Material and Methods

### Fine-mapping of the *Z*-locus in perennial ryegrass

The two perennial ryegrass populations used for fine-mapping (VrnA-XL and DTZ) were designed to segregate for the *Z*-locus as described by Manzanares et al. (2016). VrnA-XL had its origin in the VrnA population (Jensen et al. 2005), initially derived from a cross between a genotype of the Italian cultivar ‘Veyo2’ and an ecotype collected on the Danish island Falster. The F1 genotype F1-30 (SI composition S_12_Z_22_) was clonally propagated at a large scale and pollinated with pollen from a second F1 genotype (F1-39, S_12_Z_12_). The resulting offspring, i.e., seeds harvested on F1-30, were supposed to be heterozygous at the *Z*-locus. Homozygosity at the *Z*-locus either indicated rarely occurring self-pollination or a recombination event between the marker under investigation and the *Z*-locus. Similarly, DTZ originated from the perennial ryegrass ILGI mapping family (Jones et al. 2002) and was developed by crossing the ILGI siblings P150/112/129 (*S*_12_*Z*_13_) and P150/112/132 (*S*_12_*Z*_12_) but in the opposite direction as reported in Manzanares et al. (2016), i.e., P150/112/129 as male and P150/112/132 as the female parent.

Single seeds of both VrnA-XL and DTZ were grown in soil-filled plastic trays (8 × 12 pots), covered by a thin layer of sand. Around four weeks after germination, young leaf samples (approximately 15 cm long) were collected in 96-well collection plates and used for high-throughput DNA extraction as described in Manzanares et al. (2016).

For fine-mapping and BAC library screening, publicly available DNA markers from Hackauf and Wehling (2005) and Shinozuka et al. (2010) were used. Additional markers were developed by alignment of P205C9H17P and additional BAC clone sequences kindly provided by Prof. Iain Armstead, later published by Harper et al. (2019), with the rice genome sequence (RAP Build 3 *of O. sativa japonica*, NCBI) using BLASTN analysis. Primers were designed in regions being conserved between rice and perennial ryegrass using the Primer3 software (Untergasser et al. 2012). Markers were designed to amplify PCR products of 80-150 bp, suitable for high-resolution melting (HRM) analysis of unknown DNA sequence polymorphisms as described by Studer et al. (2009). Genotyping of VrnA-XL and DTZ was done at high throughput using HRM analysis as described by Manzanares et al. (2016).

### Construction of the Poaceae synteny maps

The gene annotations from the *L. perenne* genotype Kyuss (Frei et al. 2021), the *L. perenne* genotype P226/135/16 (Begheyn et al. 2018), and the *L. multiflorum* genome assembly (Copetti et al. 2021) were used to obtain the sequences of the genes within the *S*-locus and the *Z*-locus. For this purpose, the flanking markers of the *S*-locus (05_02790 and 05_02889, Manzanares et al. 2016) and the *Z*-locus (CADELP, 37600, BAC_BEG, 171R) were used to identify the *S*- and the *Z*-locus, respectively. Gene annotations were considered if an ortholog sequence was present within the *S*-locus and *Z*-locus of *L. perenne* Kyuss, *L. perenne* P226/135/16, and *L. multiflorum*; otherwise, they were removed from further analysis. If no annotation was present for an identified coding sequence, the gene structure was added using the Augustus (Organism: *O. brachyantha* L.) gene prediction tool or manually through a BLAST-based approach (Stanke and Morgenstern 2005). Furthermore, the intron-exon structure leading to the coding sequence was further streamlined for all the genes in the three genome assemblies using the open reading frame finder from NCBI combined with a BLAST-based manual approach. Therefore, the coding sequence based on the annotation file does not always perfectly align with the coding sequence used in this study. For example, within the three high-quality genome assemblies displayed in Table 1, 16 *SI-DUF247* gene sequences were identified. Of these 16 identified gene sequences, only four were annotated as an intronless gene, whereas the other sequences were either not annotated or annotated with minor or major deviations toward the intronless gene structure. All 16 gene sequences identified were then streamlined into an intronless gene structure of similar size.

Using the CLC Genomics Workbench software 11.0 (CLC bio, Aarhus, Denmark), the identified *S*- and *Z*-locus genes and flanking marker amplicon sequences were used as a query for BLAST analysis against a database containing 13 Poaceae genome assemblies (Table 2). BLAST hits were mapped if the BLAST E-value was below 1E^−80^ for all the genes except *sS* and *sZ*. For *sS* and *sZ*, the BLAST E-value was 1E^−10^. Furthermore, when a new orthologous sequence of *sS* or *sZ* was identified, it was added to the BLAST query. For the amplicon sequences of the flanking markers, the BLAST E-value was 1E^−10^. The BLAST-based annotation files of the *S*- and *Z*-locus were then translated into a CMAP file format as Veltri et al. (2016) described. The CMAP files were used to generate synteny maps using the advanced mode of the SimpleSynteny tool (Veltri et al. 2016). The graphical representation (e.g., fill colors of the structures) of the synteny maps was adopted using the Affinity Designer software (Serif (Europe) Ltd, West Bridgford, United Kingdom).

### Assessment of the functionality of the self-incompatibility (SI) candidates

The *SI-DUF247s*, the *sS*, and the *sZ* gene sequences identified were defined to be functional if they displayed a similar gene sequence and gene structure as the orthologous gene sequences found in the *L. perenne* genotypes Kyuss (Frei et al. 2021) and P226/135/16 (Byrne et al. 2015) and *L. multiflorum* (Copetti et al. 2021). Therefore an *SI-DUF247* is defined as functional if an intronless open reading frame could be found, leading to a protein size of 480 to 600 amino acids. Furthermore, the translated protein must belong to the protein family DUF247 (pfam03140) and have a predicted non-cytoplasmic domain at the C-terminus, followed by a transmembrane domain and a small cytoplasmic domain at the N-terminus according to InterProScan (Blum et al. 2021).

The *sS* and *sZ* were determined to be functional if an open reading frame with one intron could be found, leading to a protein size of 75 to 105 amino acids. In addition, a signal peptide needs to be predicted by InterProScan (Blum et al. 2021) at the C-terminus followed by a non-cytoplasmic domain.

### Phylogenetic tree construction of *S*- and *Z*-locus genes

The *S*- and *Z*-locus gene sequences from *L. perenne* Kyuss, *L. perenne* P226/135/16, *L. perenne* S23 Z and *L. multiflorum*, and the BAC clone P205 were extracted. The intron-exon structure of all *S*- and *Z*-locus genes extracted were further streamlined, leading to a comparable gene structure using the open reading frame finder from NCBI combined with a blast-based manual approach. The Lm*TPR-2* from *L. multiflorum* scf818 (*Lmu01_818G0000280*) was excluded from analysis, as it represented a distinctive duplication to the *TPR-1* and was present only once in the six *S*-locus regions analyzed. The *LpGK-2* from *L. perenne* P226/35/16 was excluded as it seemed to display a truncated duplication of the *GK-1*, and no streamlined coding sequence could be found. The coding sequence for 44 *S*-locus and 75 *Z*-locus gene sequences were extracted. The duplications of *NBS-LRR, GK*, and *PLP* were pooled. TranslatorX was used for amino acid-directed multiple sequence alignment for each group of orthologous genes (Abascal et al. 2010) by using the MAFFT algorithm v7.147b (Katoh et al. 2002). An additional alignment curation step was performed using Gblocks v0.91b with the minimal block length of five amino acids (Talavera and Castresana 2007). The alignment file was then transformed into the PHYLIP format using EasycodeML v1.2 (Gao et al. 2019). The phylogenetic trees were built using PhyML-3.1 (Guindon et al. 2010). The trees were visualized using the ggtree package in R statistical environment, version 4.1.1 (Yu 2020).

### Pairwise comparison of protein sequences of the putative SI determinants

A multiple protein sequence alignment of the *Lolium* male SI candidate genes (*SI-DUF247s*) and the *Lolium* female SI candidate genes (*sS* and *sZ*) was performed using the T-Coffee multiple sequence alignment package provided by EMBL-EBI (Madeira et al. 2019). The calculated percentage identity matrix was converted into a heat map for graphical representation.

### Expression pattern analysis of *S*- and *Z*-locus genes using RT-qPCR

#### Plant material and growth conditions

The self-incompatible and highly heterozygous *L. perenne* genotype S23 Z (Valentine and Charles 1975) was vernalized over the winter outdoors in Eschikon, Switzerland. A total of six clones were transferred into a climate chamber in the spring once the first signs of flowering (emerging of flower heads) were visible. The plants were grown under long-day conditions (16 hours light; 8 hours dark) with temperatures ranging from 20 °C during the night and 24 °C during the day.

#### Tissue sampling

Anther and stigma tissue was collected during flowering at three different time points (T_0_: one week before flowering, T_1_: two to three days before flowering, T_2_: on the day of flowering). Leaf tissue was collected as a control and did not have a specific time point (T_NA_). The sampled tissue was transferred in a 1.5 ml Eppendorf tube, immediately frozen in liquid nitrogen, and stored in a −80 °C freezer. Anther tissue was collected instead of pollen tissue as a sufficient amount of pollen for RNA extraction prior to the day of flowering (T_2_) is not possible with *L. perenne*.

#### RNA extraction and cDNA synthesis

Plastic grinding pestles were used to homogenize 45 to 80 mg of plant tissue in a 1.5 ml Eppendorf tube. The ground tissue was used for RNA extraction using the Qiagen RNeasy Mini Kit, following the “Purification of Total RNA from Plant Cells and Tissues and Filamentous Fungi” protocol (Qiagen, Hilden, Germany). Furthermore, an additional on-column DNase Digestion with the RNAse-Free DNase Set was performed according to the manufacturer’s protocol (Qiagen, Hilden, Germany). The integrity of the total RNA extracted was confirmed with the TapeStation 2200 using RNA screen tape (Agilent Technologies, Santa Clara, CA, USA). RNA samples with a RNA integrity number (RIN) value below 4.5 were discarded (Schroeder et al. 2006). The RNA quantity was determined using the Qubit BR RNA assay (Thermo Fischer Scientific, Waltham, USA). The weight, the concentration in ng/μl, and the individual RIN values can be seen in Supplementary Table 3. Double-stranded cDNA was synthesized from 0.3 μg to 1 μg of RNA using the RevertAid First Strand cDNA Synthesis Kit (Thermo Fischer Scientific, Waltham, MA, USA), following the manufacturer’s protocol with 0.5 μl Oligo (dT)18 primer and 0.5 μl Random Hexamer primers. For each RNA sample that was reverse transcript (RT sample), a no-reverse transcriptase control (NoRT sample) was included to detect a possible genomic DNA contamination of the RNA samples.

#### Primer design

Primer pairs were designed for multiple genes of interest (GOI) co-segregating with the *S*- and the *Z*-locus of perennial ryegrass (Supplementary Table 2). Furthermore, primer pairs were designed for the reference genes *EF1-α, elF4A-2, CPB20*, and *elf4A-1* as they showed a conserved expression level between pollen and stigma samples (Manzanares et al., 2016). The unresolved diploid genome assembly of *L. perenne* S23 Z was used to obtain the sequences of the GOI and the reference genes. The software Primer3 (Untergasser et al. 2012) was used to design primers leading to a product size of 75-160 bp, a primer melting temperature of 60 °C, and a primer size of 18-23 bp. The *SDUF247-I, SDUF247-II, LpsS, ZDUF247-I, ZDUF247-II*, and *LpsZ* displayed a high sequence diversity between the two alleles, making it necessary to design allele-specific primers. The sequences of the primers for the amplification of the GOI and the reference genes are displayed in Supplementary Table 2.

#### RT-qPCR data acquisition

The RT-qPCR was performed on the high-throughput BioMark HD system using a 192.24 Dynamic Array™ (Fluidigm, South San Francisco, CA, USA). All of the samples were pre-amplified for 17 cycles before the initial run, following the manufacturer’s protocol, and the final product was diluted fivefold. The samples and four negative control samples (ddH_2_0) were loaded in duplicates according to Fluidigm’s EvaGreen DNA-binding dye protocol onto the BioMark HD system. The following qPCR conditions were used: 95 °C for 30 seconds, 40 cycles of 95 °C for 5 seconds, and 60 °C for 20 seconds, plus a melting curve analysis. The data were processed using the software Fluidigm Real-Time PCR analysis 4.0 (Fluidigm, South San Francisco, CA, USA). The quality threshold was set to the default value of 0.65. Furthermore, a linear baseline correction was performed, and the setting automatic detectors were used as the C_t_ threshold method.

#### RT-qPCR data analysis

The C_t_ values were exported using the Fluidigm Real-Time PCR analysis 4.0 software (Fluidigm, South San Francisco, CA, USA). All 16 GOI and the four reference genes consistently showed a single amplicon peak. Measured C_t_ values over 21 were set to 999. This specific threshold was chosen as we observed a high standard deviation between the technical replicates with values above 21. The standard deviation between the two technical replicates for each sample was calculated, and data points with more than a 0.5 C_t_ difference were excluded from downstream analysis. Besides, only data were used where the percent deviation between the two technical replicates was below 3%. Furthermore, the four ddH_2_0 controls were assessed for each primer pair to exclude possible contamination in the chemicals. The gDNA contamination was assessed by comparing the RT samples’ C_t_ value with the noRT samples’ C_t_ value for each of the four reference genes. The gDNA contamination was considered negligible when the difference in C_t_ value between the RT sample and the noRT sample was above ten cycles. The primer efficiencies were calculated using LinReg PCR 7.5 (Ramakers et al. 2003) and are shown in Supplementary Table 2. The stability of the reference genes was assessed using GeNorm (Huang et al. 2014). The expression stability value for *EF1-alpha* was 0.292, for *CPB20* 0.488, for *eIF4A-2* 0.292, and for *eIF4A-1* 0.385, meaning all four reference genes qualify to be used as they all have an expression stability value below 1.5 which represent the geNorm cut off (Vandesompele et al. 2002).

To determine the relative gene expression ratio of the GOI, a relative quantification method as described by Pfaffl (2001) was used. The ΔC_t_ value was calculated by subtracting the C_t_ value of the sample minus the C_t_ value of the control. *LpTIR, LpdsRNAbp, SNF2, LpGK, LpSDUF247-I, LpSDUF247-II, LpZDUF247-I*, and *LpZDUF247-II*, the first biological replicate of the anther tissue at time point 1 was used as the control. For *LpsS* and *LpsZ*, the first biological replicate of the stigma tissue at time point 1 was used as the control. The ΔC_t_ was set to the power of the respective PCR efficiency, leading to the relative quantity (RQ) value. The relative gene expression was calculated by dividing the RQ value of the GOI by the geometrical mean RQ value of the reference gene. The relative gene expression ratios of the *S*- and *Z*-locus genes were displayed in polar charts, and a scatter plot using the R statistical environment, version 4.1.1.

In addition, the ΔC_t_ values of *S*- and *Z*-locus genes were calculated to compare the expression levels of different genes in the same sample. The ΔC_t_ values were calculated as the C_t_ value of GOI minus the geometrical mean C_t_ value of the four reference genes (*EF1-alpha, CPB20, eIF4A-2, eIF4A-1*). The ΔC_t_ values were then displayed in a heat map using the R statistical environment, version 4.1.1. No scaling was applied, and the data were clustered on the level of genes (rows).

## Availability of data and materials

The data produced for this project has been deposited in the NCBI database under the BioProject NNNNNNN. The sequence of the BAC clone P205C9H17P is available under the accession NNNNNNNN. The scaffolds spanning the *S*- and the *Z*-locus from *L. perenne* S23 Z and *L. multiflorum* are available under the accession NNNNNNNN. The coding sequence of all genes identified within the *S*- and *Z*-locus within *Lolium* species are deposited under the accession NNNNNNNN.

## Acknowledgments

We thank Stephan Hentrup at Aarhus University for plant material development and maintenance, Dr Zeljko Micic from Deutsche Saatveredelung AG for helpful advice in the lab, and Dr. Maurice Bosch (IBERS, Aberystwyth) for providing us with the perennial ryegrass genotype S23 Z. The RT-qPCR data was generated at the Genetic Diversity Centre (GDC) of ETH Zurich. For assistance in the laboratory with the RNA extraction, reverse transcription, and expression data acquisition and analysis using BioMark HD, we thank Silvia Kobel (GDC, ETH Zurich), Dr. Aria Minder (GDC, ETH Zurich), and Dr. Niklaus Zemp (GDC, ETH Zurich). Further, we want to thank Verena Knorst-Rashid (Molecular Plant Breeding, ETH Zurich) for her assistance in maintaining the *L. perenne* S23 plants as well as Prof. Dr. Achim Walter (Crop Science, ETH Zurich) for allowing us access to the greenhouse and laboratory infrastructures.

This study was funded by the Swiss National Science Foundation (SNSF, project number 310030_197708), the SNSF professorship with the grant number PP00P2_138983 and supported by the Danish Council for Independent Research, Technology and Production Sciences, Denmark.

