## Supplementary Figure 1 to 3 and Supplementary Table 1 to 3 for "Fine-mapping and comparative genomic analysis reveal the gene composition at the *S* and *Z* self-incompatibility loci in grasses"

### Supplementary material

**A**

| | Species and genotype | Localization | Protein name | 1. | 2. | 3. | 4. | 5. | 6. | 7. | 8. | 9. | 10. | 11. | 12. | 13. | 14. | 15. | 16. | 17. | 18. | 19. | 20. | 21. | 22. | 23. | 24. | 25. | 26. |
|---|---|---|---|---|---|---|---|---|---|---|---|---|---|---|---|---|---|---|---|---|---|---|---|---|---|---|---|---|---|
| | | | | **SDUF247-I** | | | | | | **SDUF247-II** | | | | | | **ZDUF247-I** | | | | | | | **ZDUF247-II** | | | | | | |
| **1.** | *L. perenne* Kyuss | chr1 | **SDUF247-I** | 100 | 80.72 | 78.81 | 81.85 | 78.81 | 77.36 | 43.02 | 42.77 | 41.01 | 43.53 | 41.01 | 42.26 | 45.69 | 43.46 | 42.88 | 43.46 | 42.91 | 44.38 | 42.52 | 41.38 | 41.84 | 41.15 | 41.84 | 38.96 | 41.17 | 40.73 |
| **2.** | *L. perenne* P226/135/16 | chr1 | | 80.72 | 100 | 77.23 | 81.36 | 77.23 | 79.85 | 42.75 | 42.5 | 41.31 | 44.23 | 41.31 | 42.94 | 46.97 | 43.19 | 43.57 | 43.19 | 42.64 | 44.29 | 42.64 | 40.54 | 41.57 | 41.35 | 41.57 | 38.12 | 39.15 | 40.66 |
| **3.** | *L. perenne* S23 Z | contig11029 | | 78.81 | 77.23 | 100 | 78.02 | 100 | 82.61 | 43.5 | 43.43 | 42.6 | 43.9 | 42.6 | 44.49 | 43.82 | 42.38 | 42.97 | 42.38 | 43.4 | 43.71 | 42.54 | 41.78 | 42.25 | 41.65 | 42.25 | 40.04 | 41.33 | 42 |
| **4.** | *L. perenne* S23 Z | contig12948 | | 81.85 | 81.36 | 78.02 | 100 | 78.02 | 79.1 | 44.1 | 44.23 | 42.47 | 45.4 | 42.47 | 44.08 | 46.77 | 44.72 | 44.15 | 44.72 | 43.6 | 44.68 | 43.8 | 41.49 | 42.72 | 40.96 | 42.72 | 38.89 | 41.28 | 41.23 |
| **5.** | *L. multiflorum* | scf1212 | | 78.81 | 77.23 | 100 | 78.02 | 100 | 82.61 | 43.5 | 43.43 | 42.6 | 43.9 | 42.6 | 44.49 | 43.82 | 42.38 | 42.97 | 42.38 | 43.4 | 43.71 | 42.54 | 41.78 | 42.25 | 41.65 | 42.25 | 40.04 | 41.33 | 42 |
| **6.** | *L. multiflorum* | scf818 | | 77.36 | 79.85 | 82.61 | 79.1 | 82.61 | 100 | 42.08 | 42.42 | 40.85 | 43.16 | 40.85 | 42.48 | 45.31 | 43.3 | 43.49 | 43.3 | 41.59 | 42.47 | 42.17 | 39.31 | 40.15 | 39.35 | 40.15 | 38.05 | 40.04 | 39.62 |
| **7.** | *L. perenne* Kyuss | chr1 | **SDUF247-II** | 43.02 | 42.75 | 43.5 | 44.1 | 43.5 | 42.08 | 100 | 76.18 | 82.18 | 75.85 | 82.18 | 72.45 | 46.84 | 43.41 | 42.83 | 43.41 | 45.42 | 45.61 | 46.86 | 38.54 | 42.69 | 42.47 | 42.69 | 41.15 | 40.96 | 41.36 |
| **8.** | *L. perenne* P226/135/16 | chr1 | | 42.77 | 42.5 | 43.43 | 44.23 | 43.43 | 42.42 | 76.18 | 100 | 74.59 | 76.23 | 74.59 | 72.5 | 47.64 | 44.51 | 44.89 | 44.51 | 46.71 | 46.71 | 46.98 | 40.04 | 43.68 | 42.12 | 43.68 | 41.49 | 41.11 | 41.31 |
| **9.** | *L. perenne* S23 Z | contig11029 | | 41.01 | 41.31 | 42.6 | 42.47 | 42.6 | 40.85 | 82.18 | 74.59 | 100 | 74.91 | 100 | 71.58 | 47.04 | 43.13 | 42.55 | 43.13 | 46.11 | 45.72 | 46.97 | 37.5 | 41.92 | 40.73 | 41.92 | 42.03 | 42.03 | 41.67 |
| **10.** | *L. perenne* S23 Z | contig11774 | | 43.53 | 44.23 | 43.9 | 45.4 | 43.9 | 43.16 | 75.85 | 76.23 | 74.91 | 100 | 74.91 | 72.97 | 47.52 | 44.81 | 42.86 | 44.81 | 45.67 | 46.26 | 47.14 | 40.08 | 43.77 | 42.38 | 43.77 | 42.41 | 42.02 | 41.76 |
| **11.** | *L. multiflorum* | scf1212 | | 41.01 | 41.31 | 42.6 | 42.47 | 42.6 | 40.85 | 82.18 | 74.59 | 100 | 74.91 | 100 | 71.58 | 47.04 | 43.13 | 42.55 | 43.13 | 46.11 | 45.72 | 46.97 | 37.5 | 41.92 | 40.73 | 41.92 | 42.03 | 42.03 | 41.67 |
| **12.** | *L. multiflorum* | scf818 | | 42.26 | 42.94 | 44.49 | 44.08 | 44.49 | 42.48 | 72.45 | 72.5 | 71.58 | 72.97 | 71.58 | 100 | 47.08 | 42.48 | 42.48 | 42.48 | 45.4 | 46.55 | 45.66 | 39.28 | 42.61 | 40.87 | 42.61 | 40.72 | 42.45 | 41.49 |
| **13.** | *L. perenne* Kyuss | chr2 | **ZDUF247-I** | 45.69 | 46.97 | 43.82 | 46.77 | 43.82 | 45.31 | 46.84 | 47.64 | 47.04 | 47.52 | 47.04 | 47.08 | 100 | 58.22 | 57.84 | 58.22 | 57.25 | 58.75 | 59.36 | 45.16 | 47.1 | 48.16 | 47.1 | 46.36 | 45.29 | 45.02 |
| **14.** | *L. perenne* F1-30 | P205 | | 43.46 | 43.19 | 42.38 | 44.72 | 42.38 | 43.3 | 43.41 | 44.51 | 43.13 | 44.81 | 43.13 | 42.48 | 58.22 | 100 | 73.62 | 100 | 55.7 | 56.24 | 58.61 | 42.3 | 43.29 | 43.73 | 43.29 | 43.71 | 42.42 | 43.9 |
| **15.** | *L. perenne* P226/135/16 | chr2 | | 42.88 | 43.57 | 42.97 | 44.15 | 42.97 | 43.49 | 42.83 | 44.89 | 42.55 | 42.86 | 42.55 | 42.48 | 57.84 | 73.62 | 100 | 73.62 | 54.02 | 54.38 | 57.68 | 43.41 | 43.1 | 43.54 | 43.1 | 41.09 | 41.65 | 44.09 |
| **16.** | *L. perenne* S23 Z | contig55097 | | 43.46 | 43.19 | 42.38 | 44.72 | 42.38 | 43.3 | 43.41 | 44.51 | 43.13 | 44.81 | 43.13 | 42.48 | 58.22 | 100 | 73.62 | 100 | 55.7 | 56.24 | 58.61 | 42.3 | 43.29 | 43.73 | 43.29 | 43.71 | 42.42 | 43.9 |
| **17.** | *L. perenne* S23 Z | contig7728 | | 42.91 | 42.64 | 43.4 | 43.6 | 43.4 | 41.59 | 45.42 | 46.71 | 46.11 | 45.67 | 46.11 | 45.4 | 57.25 | 55.7 | 54.02 | 55.7 | 100 | 79 | 57.47 | 42.29 | 44.57 | 44.25 | 44.57 | 44.4 | 43.13 | 43.07 |
| **18.** | *L. multiflorum* | scf1905 | | 44.38 | 44.29 | 43.71 | 44.68 | 43.71 | 42.47 | 45.61 | 46.71 | 45.72 | 46.26 | 45.72 | 46.55 | 58.75 | 56.24 | 54.38 | 56.24 | 79 | 100 | 58.76 | 42.51 | 45.44 | 44.36 | 45.44 | 43.86 | 43.71 | 44.61 |
| **19.** | *L. multiflorum* | scf3448 | | 42.52 | 42.64 | 42.54 | 43.8 | 42.54 | 42.17 | 46.86 | 46.98 | 46.97 | 47.14 | 46.97 | 45.66 | 59.36 | 58.61 | 57.68 | 58.61 | 57.47 | 58.76 | 100 | 42.29 | 42.53 | 44.32 | 42.53 | 42.69 | 43.5 | 43.07 |
| **20.** | *L. perenne* Kyuss | chr2 | **ZDUF247-II** | 41.38 | 40.54 | 41.78 | 41.49 | 41.78 | 39.31 | 38.54 | 40.04 | 37.5 | 40.08 | 37.5 | 39.28 | 45.16 | 42.3 | 43.41 | 42.3 | 42.29 | 42.51 | 42.29 | 100 | 49.43 | 50.47 | 49.43 | 50.28 | 50.47 | 51.39 |
| **21.** | *L. perenne* F1-30 | P205 | | 41.84 | 41.57 | 42.25 | 42.72 | 42.25 | 40.15 | 42.69 | 43.68 | 41.92 | 43.77 | 41.92 | 42.61 | 47.1 | 43.29 | 43.1 | 43.29 | 44.57 | 45.44 | 42.53 | 49.43 | 100 | 67.04 | 100 | 50.28 | 51.06 | 50.86 |
| **22.** | *L. perenne* P226/135/16 | chr2 | | 41.15 | 41.35 | 41.65 | 40.96 | 41.65 | 39.35 | 42.47 | 42.12 | 40.73 | 42.38 | 40.73 | 40.87 | 48.16 | 43.73 | 43.54 | 43.73 | 44.25 | 44.36 | 44.32 | 50.47 | 67.04 | 100 | 67.04 | 49.81 | 47.97 | 50.67 |
| **23.** | *L. perenne* S23 Z | contig55097 | | 41.84 | 41.57 | 42.25 | 42.72 | 42.25 | 40.15 | 42.69 | 43.68 | 41.92 | 43.77 | 41.92 | 42.61 | 47.1 | 43.29 | 43.1 | 43.29 | 44.57 | 45.44 | 42.53 | 49.43 | 100 | 67.04 | 100 | 50.28 | 51.06 | 50.86 |
| **24.** | *L. perenne* S23 Z | contig4538 | | 38.96 | 38.12 | 40.04 | 38.89 | 40.04 | 38.05 | 41.15 | 41.49 | 42.03 | 42.41 | 42.03 | 40.72 | 46.36 | 43.71 | 41.09 | 43.71 | 44.4 | 43.86 | 42.69 | 50.28 | 50.28 | 49.81 | 50.28 | 100 | 73.5 | 49.44 |
| **25.** | *L. multiflorum* | scf1905 | | 41.17 | 39.15 | 41.33 | 41.28 | 41.33 | 40.04 | 40.96 | 41.11 | 42.03 | 42.02 | 42.03 | 42.45 | 45.29 | 42.42 | 41.65 | 42.42 | 43.13 | 43.71 | 43.5 | 50.47 | 51.06 | 47.97 | 51.06 | 73.5 | 100 | 48.76 |
| **26.** | *L. multiflorum* | scf3448 | | 40.73 | 40.66 | 42 | 41.23 | 42 | 39.62 | 41.36 | 41.31 | 41.67 | 41.76 | 41.67 | 41.49 | 45.02 | 43.9 | 44.09 | 43.9 | 43.07 | 44.61 | 43.07 | 51.39 | 50.86 | 50.67 | 50.86 | 49.44 | 48.76 | 100 |

**B**

| | Species and genotype | Localization | Protein name | 1. | 2. | 3. | 4. | 5. | 6. | 7. | 8. | 9. | 10. | 11. | 12. | 13. |
|---|---|---|---|---|---|---|---|---|---|---|---|---|---|---|---|---|
| | | | | **sS** | | | | | | **sZ** | | | | | | |
| **1.** | *L. perenne* Kyuss | chr1 | **sS** | 100 | 52.44 | 62.65 | 48.15 | 62.65 | 57.32 | 30.16 | 30.16 | 27.94 | 20 | 30.16 | 22.95 | 24.62 |
| **2.** | *L. perenne* P226/135/16 | chr1 | | 52.44 | 100 | 53.57 | 55 | 53.57 | 60.71 | 35.29 | 30.88 | 31.51 | 21.54 | 30.88 | 24.62 | 24.29 |
| **3.** | *L. perenne* S23 Z | contig11029 | | 62.65 | 53.57 | 100 | 39.51 | 100 | 51.19 | 29.23 | 30.77 | 26.76 | 22.58 | 30.77 | 24.19 | 19.4 |
| **4.** | *L. perenne* S23 Z | contig12948 | | 48.15 | 55 | 39.51 | 100 | 39.51 | 53.09 | 30.65 | 32.26 | 37.31 | 33.9 | 32.26 | 33.33 | 32.81 |
| **5.** | *L. multiflorum* | scf1212 | | 62.65 | 53.57 | 100 | 39.51 | 100 | 51.19 | 29.23 | 30.77 | 26.76 | 22.58 | 30.77 | 24.19 | 19.4 |
| **6.** | *L. multiflorum* | scf818 | | 57.32 | 60.71 | 51.19 | 53.09 | 51.19 | 100 | 27.69 | 29.23 | 32.86 | 25.81 | 29.23 | 24.19 | 26.87 |
| **7.** | *L. perenne* Kyuss | chr2 | **sZ** | 30.16 | 35.29 | 29.23 | 30.65 | 29.23 | 27.69 | 100 | 31.08 | 34.18 | 35.62 | 31.08 | 32.43 | 29.73 |
| **8.** | *L. perenne* F1-30 | P205 | | 30.16 | 30.88 | 30.77 | 32.26 | 30.77 | 29.23 | 31.08 | 100 | 53.26 | 24.39 | 100 | 24.39 | 28.05 |
| **9.** | *L. perenne* P226/135/16 | chr2 | | 27.94 | 31.51 | 26.76 | 37.31 | 26.76 | 32.86 | 34.18 | 53.26 | 100 | 28.41 | 53.26 | 21.84 | 34.52 |
| **10.** | *L. perenne* S23 Z | contig4358 | | 20 | 21.54 | 22.58 | 33.9 | 22.58 | 25.81 | 35.62 | 24.39 | 28.41 | 100 | 24.39 | 53.75 | 30.26 |
| **11.** | *L. perenne* S23 Z | contig55097 | | 30.16 | 30.88 | 30.77 | 32.26 | 30.77 | 29.23 | 31.08 | 100 | 53.26 | 24.39 | 100 | 24.39 | 28.05 |
| **12.** | *L. multiflorum* | scf1905 | | 22.95 | 24.62 | 24.19 | 33.33 | 24.19 | 24.19 | 32.43 | 24.39 | 21.84 | 53.75 | 24.39 | 100 | 26.32 |
| **13.** | *L. multiflorum* | scf3448 | | 24.62 | 24.29 | 19.4 | 32.81 | 19.4 | 26.87 | 29.73 | 28.05 | 34.52 | 30.26 | 28.05 | 26.32 | 100 |

**C**

| |
|---|
| 100 |
| 90 |
| 80 |
| 70 |
| 60 |
| 50 |
| 40 |
| 30 |
| 20 |

Supplementary Figure 1: Pairwise comparison of the protein sequences of the male and female self-incompatibility candidates. (A) Pairwise comparison for the SI-DUF247s. (B) Pairwise comparison for the sS and sZ. (C) Color code indicating the protein sequence identity in percentage.

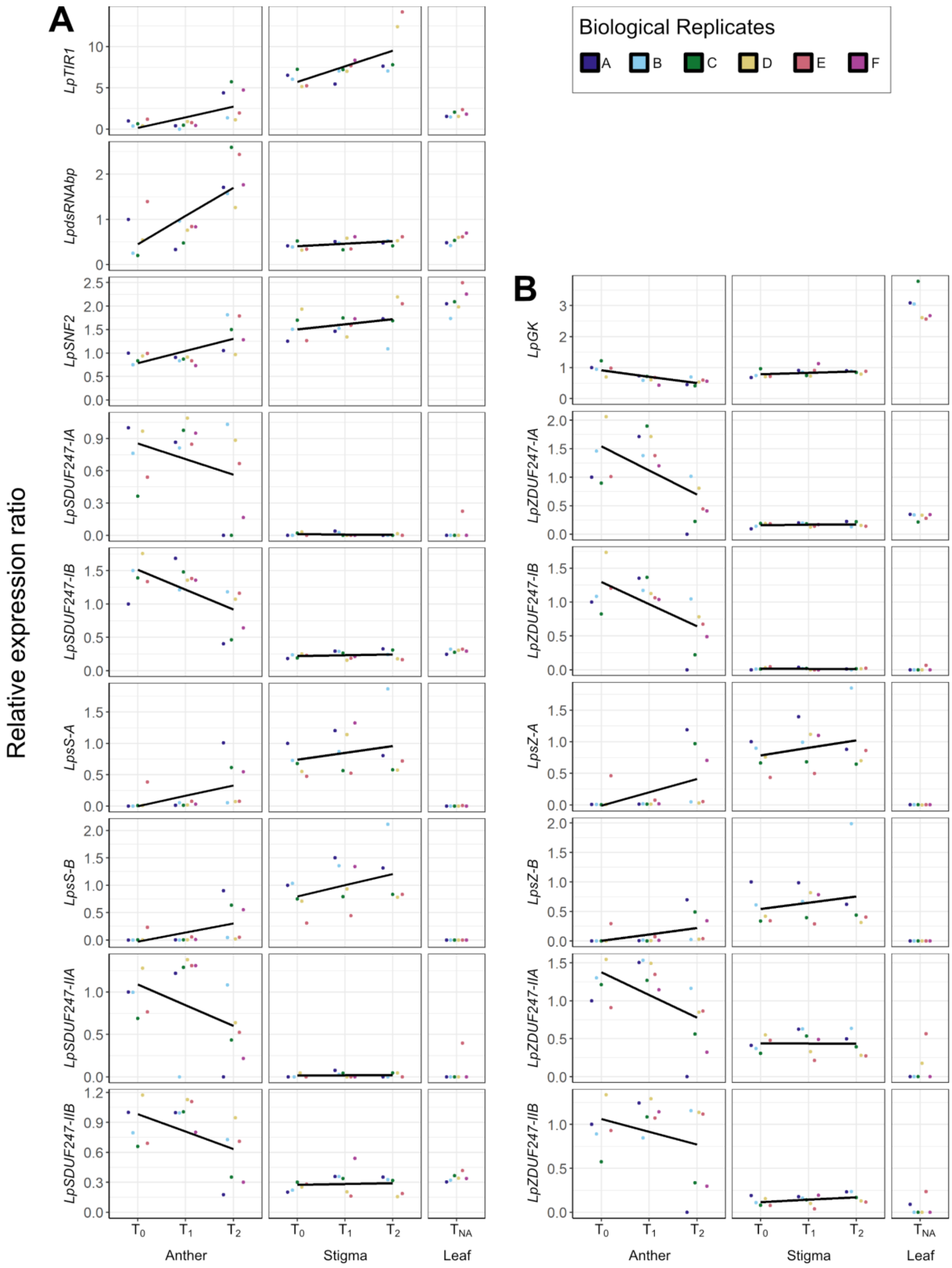


Supplementary Figure 2: Relative expression ratio of *S*- and *Z*-locus genes in *Lolium perenne* L. genotype S23 Z. The relative expression ratios were calculated according to Pfaffl (2001) and normalized to four reference genes (*EF1-alpha*, *CPB20*, *eIF4A-2*, *eIF4A-1*). For *LpsS* and *LpsZ*, the expression value in stigma tissue at time point 0 ($T_0$) in the first biological replicate was set as 1. For all the other genes displayed, the expression value in anthers at time point 0 ($T_O$) in the first biological replicate was set to 1. The color scheme in the top right corner is used to distinguish the biological replicates. The relative expression ratios can be seen on the Y-axis and the different sample tissues at different time points on the X-axis. Furthermore, for the *LpsS*, *LpsZ*, and the *Sl-DUF247*s, the relative expression ratio measurement was explicitly performed for each allele present in the genotype. The two different alleles are indicated by A and B after the gene name. Only data points were included, where the $C_t$ difference was below 0.5 and the percent deviation was below 3% between the two technical replicates. Therefore, the standard error is not displayed in the figure. (A) Relative expression ratios of *S*-locus genes. (B) Relative expression ratios of *Z*-locus genes.

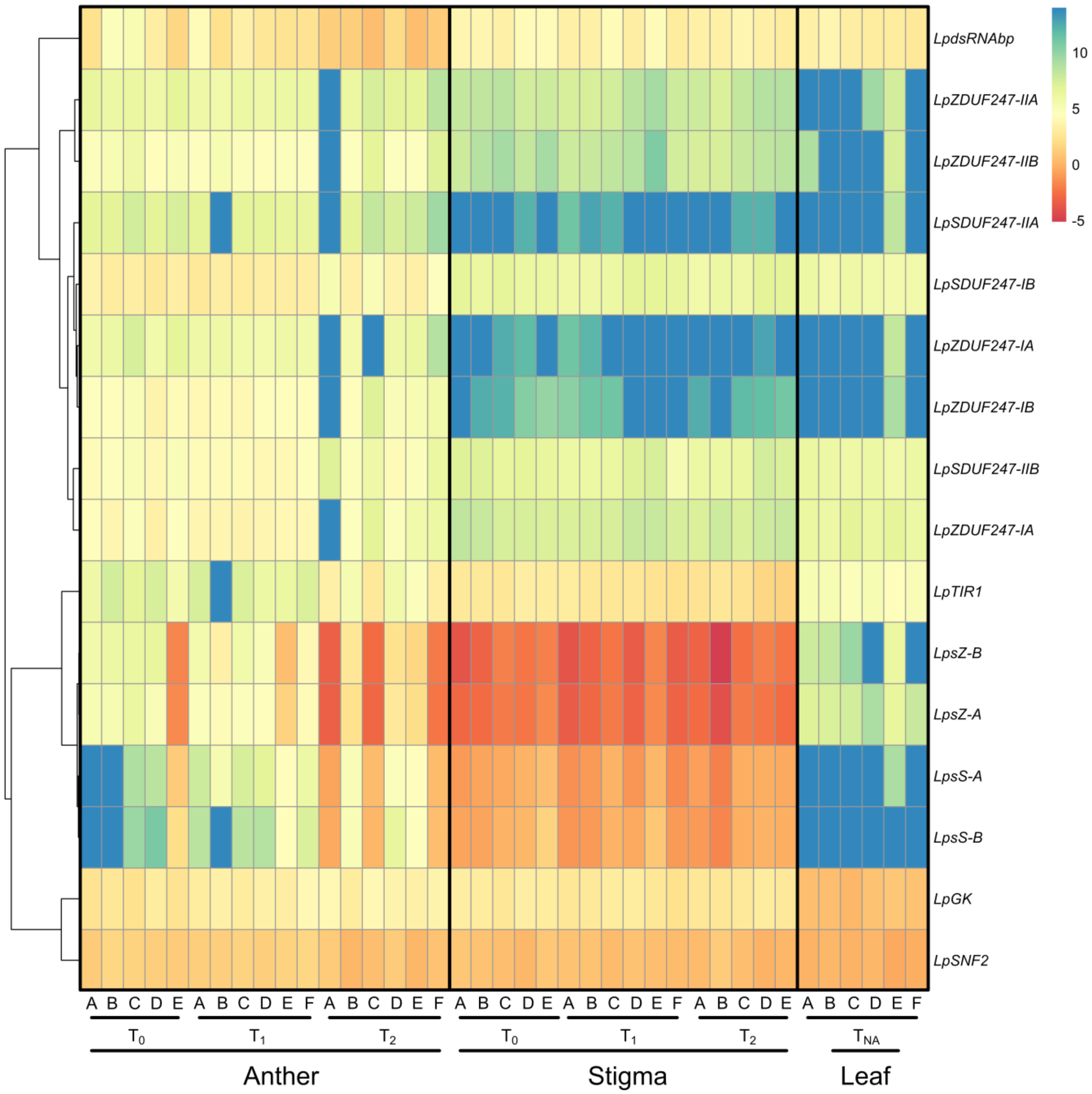


Supplementary Figure 3: Expression levels of *S*- and *Z*-locus genes relative to four reference genes in *Lolium perenne* L. genotype S23 Z. The $\Delta C_t$ values were calculated as the $C_t$ value of the gene of interest minus the geometrical mean $C_t$ value of the four reference genes (*EF1-alpha, CPB20, eIF4A-2, eIF4A-1)*. The rows indicate the different genes localized within the *S* and *Z* loci. For the *LpsS*, *LpsZ* and the *SI-DUF247*s, the $\Delta C_t$ was calculated for both alleles present in the genotype. The two different alleles are indicated by A and B after the gene name. The columns indicate the different sample types at different time points. The biological replicates are indicated with A-F. No scaling was applied, and the data were clustered on the level of genes (rows). A dendrogram is drawn on the left, illustrating the hierarchical relationship between the expression pattern of the genes. Blue corresponds to low expression, whereas red corresponds to high expression.

Supplementary Table 1: Number of the recombination events observed between molecular markers at the *Z*-locus region in the VrnA-XL and DTZ population.

| Marker names | Forward primer sequence | Reverse primer sequence | VrnA-XL | DTZ |
|---|---|---|---|---|
| CADELP | TGCTCTTGAGGTTTGGCTAT | ATTGAGTCTTCAACAGCATTTTT | 22 | 48 |
| 37600 | AAGTGGGACAAAAGGGAACC | ATGAAGTACAAGCAAGAACAAATCA | 1 | 0 |
| 19060 | CCGATTGGTAAAAACGAGTTG | GGCAAAAGGTCAGCTTGCTA | 0 | NA [a] |
| 15550 | TGAGGTGCCTTCTGAAACTG | CGTTAGCTGGTGCTGTGGT | 0 | NA [a] |
| 12000 | CCTGGTACAAGGCGGTTTC | CGGCTAACTGATACACCACCT | 0 | NA [a] |
| 10280 | TGCCATAGTGTGGATGGATG | GCTCACTTTCCAGTCTCCTGA | 0 | 0 |
| 7800 | CGTTGATGGTGGTGCTACTG | AACGTAAAGCTAACCACAAACAGA | 0 | 0 |
| 6500 | TTTGTTTGACGCCCCTTAAC | TTCAAACCAGCATCCACATT | 0 | NA [a] |
| 5000 | CGTGCCGGAGTAGATAGAGG | GAGAGGTTATAGATGGATCCGAT | 0 | NA [a] |
| BAC_BEG | AAGCGACAATGGAAGATTCG | CAGTGGAGTTCTCCCGGTTA | 1 | 7 |
| 171R | GCGGGCGTTTCCAACTATTA | TGAATTGCACGGTAGTCCTG | 7 | NA [a] |
| Lp02_555 | GTCAGAGCTCGCACCCTTC | CCCTTGGTGATCATTGCAC | 67 | 51 |

Note: The populations have been screened for recombination using twelve markers, mapping within the *Z*-locus region. The fine-mapped *Z*-locus is defined by the region without any recombination events. In addition, the primer sequences of the polymorphic molecular markers used for the fine-mapping of the self-incompatibility locus *Z* are given.

[a] Markers were either monomorphic or did not amplify in DTZ

Supplementary Table 2: Primers used to investigate the expression pattern of *S*- and *Z*-locus genes using RT-qPCR.

| | Gene name | Primer F | Primer R | Amplicon size | Primer efficiency |
|---|---|---|---|---|---|
| ***S*-locus genes of interest** | *LpTIR1* | 5'-CAATGTTGCTAGCCTGCTTC-3' | 5'-AATGCTGGCAGGTAATCTGG-3' | 176 | 1.79 |
| | *LpdsRNA* | 5'-AGAGTCGAAAAAGCCAGCAG-3' | 5'-CTTGGTGGAGTTTGCATGTC-3' | 180 | 1.76 |
| | *LpSNF2* | 5'-TGACCCATAAACACTGTCACG-3' | 5'-CCCCATCATCTCATCCTAGTG-3' | 184 | 1.81 |
| | *LpSDUF247-IA* | 5'-GAATGCGGTTCCAGAAAAGG-3' | 5'-GCAGTGATGACATTCCCAAC-3' | 181 | 1.77 |
| | *LpSDUF247-IB* | 5'-TCTGGGCTTGAGAAAGAAGC-3' | 5'-CCTCTCGTCTACCATGTTTTCC-3' | 180 | 1.82 |
| | *LpSDUF247-IIA* | 5'-AGAAACGAGCAAGGATGTGC-3' | 5'-CTCGTGTATTTCGGCATGTG-3' | 176 | 1.91 |
| | *LpSDUF247-IIB* | 5'-TGAGGACCATTCGGCTATTG-3' | 5'-TAGCACCGTCCAATCTGTTG-3' | 177 | 1.81 |
| | *LpsS-A* | 5'-CGTCATGGTTCTGATGGTTATG-3' | 5'-GATGCCCAAATGGTTTTCC-3' | 72 | 1.78 |
| | *LpsS-B* | 5'-CGGTTTCTCCGAATCTAAGG-3' | 5'-CCTTGTACATGCAAGGTGGTC-3' | 80 | 1.66 |
| ***Z*-locus genes of interest** | *LpZDUF247-IA* | 5'-GGTGGCAATTGAGGAACTTG-3' | 5'-CTCAAGAGTGCGCATTTCAC-3' | 184 | 1.87 |
| | *LpZDUF247-IB* | 5'-GCGCAGAGTTGCACAATATC-3' | 5'-CTCGAACGCAATAAGGTTCC-3' | 175 | 1.79 |
| | *LpZDUF247-IIA* | 5'-GCGAGCAGTAGAATACCATCG-3' | 5'-GGATCTGTTTGCTCAAGTGC-3' | 183 | 1.77 |
| | *LpZDUF247B-IIB* | 5'-TGTCAGGCTAACCAAAGACG-3' | 5'-GATGACCGCAGCAAATACAG-3' | 176 | 1.80 |
| | *LpGK* | 5'-AACTGGTGCCTTCATCCTTC-3' | 5'-TAATGATCCCAAGGCTGTCC-3' | 179 | 1.77 |
| | *LpsZ-A* | 5'-TTACCCCCAACACTGAACAC-3' | 5'-AACCAGGCTGTTTGGTATGG-3' | 145 | 1.79 |
| | *LpsZ-B* | 5'-AACAAAGTGGTCGTCCTTCTG-3' | 5'-GAGAAAGTGAATCCCCATGC-3' | 97 | 1.78 |
| **Reference genes** | *EF1-alpha* | 5'-TTGACAAGCGTGTGATCGAG-3' | 5'-TGACCAGGAGCATCAATGAC-3' | 184 | 1.81 |
| | *eIF4A-2* | 5'-ATGCATGTGTTGGAGGAACC-3' | 5'-CCTTGAAACCACGAGAAAGC-3' | 183 | 1.80 |
| | *CPB20* | 5'-ACGCAGGAGGAGTTCGAG-3' | 5'-GGTCTTGGTGTTCTTGTCGAG-3' | 155 | 1.81 |
| | *eIF4A-1* | 5'-GAAAGACTGCCACCTTCTGC-3' | 5'-CGAACAGATGTTCCACCAAC-3' | 181 | 1.74 |

Note: The suffixes A and B in the gene name represent the two alleles of a specific gene. The displayed primer efficiencies were calculated using LinReg PCR 7.5 (Ramakers et al. 2003)

Supplementary Table 3: Quality and quantity assessment of the RNA samples used for RT-qPCR.

| Tissue Type | Name | Weight [g] | Conc. Qbit [ng/µl] | RIN Value |
|---|---|---|---|---|
| **Anther** | Anther $T_0$ A | 0.048 | 210 | 6.9 |
| | Anther $T_0$ B | 0.05 | 262 | 7.5 |
| | Anther $T_0$ C | 0.072 | 404 | 8.4 |
| | Anther $T_0$ D | 0.05 | 222 | 8.6 |
| | Anther $T_0$ E | 0.056 | 165 | 7 |
| | Anther $T_1$ A | 0.059 | 250 | 7.6 |
| | Anther $T_1$ B | 0.053 | 160 | 8.4 |
| | Anther $T_1$ C | 0.051 | 256 | 6 |
| | Anther $T_1$ D | 0.07 | 166 | 7.5 |
| | Anther $T_1$ E | 0.057 | 41 | 8.8 |
| | Anther $T_1$ F | 0.057 | 97 | 8.7 |
| | Anther $T_2$ A | 0.048 | 32 | 6.5 |
| | Anther $T_2$ B | 0.068 | 70 | NA |
| | Anther $T_2$ C | 0.074 | 50 | 7 |
| | Anther $T_2$ D | 0.053 | 60 | 8 |
| | Anther $T_2$ E | 0.066 | 52 | 8 |
| | Anther $T_2$ F | 0.066 | 54 | 8.4 |
| **Stigma** | Stigma $T_0$ A | 0.051 | 80 | 9.1 |
| | Stigma $T_0$ B | 0.055 | 62 | 9.3 |
| | Stigma $T_0$ C | 0.05 | 260 | 7.6 |
| | Stigma $T_0$ D | 0.055 | 78 | 9.1 |
| | Stigma $T_0$ E | 0.072 | 83 | 8.9 |
| | Stigma $T_1$ A | 0.04 | 50 | 7.6 |
| | Stigma $T_1$ B | 0.043 | 51 | 7.8 |
| | Stigma $T_1$ C | 0.045 | 42 | 5.6 |
| | Stigma $T_1$ D | 0.048 | 130 | 8.7 |
| | Stigma $T_1$ E | 0.044 | 74 | 9.5 |
| | Stigma $T_1$ F | 0.029 | 238 | 8.8 |
| | Stigma $T_2$ A | 0.047 | 122 | 9.1 |
| | Stigma $T_2$ B | 0.048 | 55 | 9.1 |
| | Stigma $T_2$ C | 0.065 | 120 | 9.1 |
| | Stigma $T_2$ D | 0.046 | 90 | 8.8 |
| | Stigma $T_2$ E | 0.051 | 129 | 9.1 |
| **Leaf** | Leaf $T_{NA}$ A | 0.041 | 47 | 5.2 |
| | Leaf $T_{NA}$ B | 0.054 | 31 | 4.5 |
| | Leaf $T_{NA}$ C | 0.064 | 33 | 5 |
| | Leaf $T_{NA}$ D | 0.056 | 117 | 5.5 |
| | Leaf $T_{NA}$ E | 0.047 | 60 | 5.6 |
| | Leaf $T_{NA}$ F | 0.046 | 51 | 5.3 |
